# Dynorphin / kappa-opioid receptor regulation of excitation-inhibition balance toggles afferent control of prefrontal cortical circuits in a pathway-specific manner

**DOI:** 10.1101/2022.10.20.513075

**Authors:** Hector E. Yarur, Sanne M. Casello, Valerie Tsai, Juan Enriquez-Traba, Rufina Kore, Huikun Wang, Miguel Arenivar, Hugo A. Tejeda

**Affiliations:** Unit on Neuromodulation and Synaptic Integration, National Institute of Mental Health, National Institutes of Health, Bethesda, MD, USA; NIH Graduate Partnership Program

## Abstract

The medial prefrontal cortex (mPFC) controls emotional behaviors and cognition via connections with limbic excitatory afferents that engage various intra-mPFC inhibitory motifs The mPFC dynorphin (Dyn) / kappa-opioid receptor (KOR) system regulates affect and cognition and is implicated in neuropsychiatric disorders. However, it’s unclear how neuropeptides in the mPFC, including the Dyn / KOR system, control excitatory and inhibitory circuit motifs integral in information processing. Here, we provide a circuit-based framework wherein selective KOR expression in mPFC afferents or within mPFC feedforward and feedback inhibitory circuits gates how distinct limbic afferent inputs control mPFC neurons. Dyn/KOR signaling directly decreases the ability of KOR-expressing afferent inputs to drive mPFC cell activity. Dyn/KOR signaling also suppresses afferent-driven recruitment of inhibitory sub-networks via several mechanisms, disinhibiting KOR-negative excitatory afferent control of mPFC ensembles. Thus, the Dyn/KOR system toggles which afferent input controls mPFC circuits, providing mechanistic insights into the role of neuropeptides in shaping mPFC function.

**Highlight:**

- Pathway-specific KOR expression confers selective filtering of mPFC afferents by dynorphin
- Endogenous dynorphin release gates KOR-expressing inputs to both dynorphin-expressing and lacking mPFC neurons
- Dynorphin / KOR modulation reveals parallel channels within amygdalo-cortical and cortical-cortical circuits
- Dynorphin disinhibits mPFC pyramidal neurons via KOR-mediated suppression of distinct inhibitory circuit motifs that preferentially impact SST-mediated feedforward inhibition
- Dynorphin / KOR signaling biases afferent control of mPFC principal cells away from KOR-positive and towards KOR-negative afferent inputs

## Introduction

Cortical circuits, including the medial prefrontal cortex (mPFC), integrate excitatory inputs with local circuit inhibition to control downstream cortical and sub-cortical targets. This function is essential for the mPFC to regulate a plethora of higher-order behaviors including action selection in response to threats/stressors or homeostatic challenges, affective behavior, motivation, and goal-directed behavior (Wassum, 2022; Alexandra Kredlow et al., 2022; Murray and Fellows, 2022; Laubach et al., 2015). Recent advances have been made in understanding the cell types and connections with limbic regions that are essential for the many facets of the mPFC in regulating behavior and the nuanced connectivity of long-range and local circuit circuits innervating and arising from within the mPFC, respectively. However, our understanding of how mPFC circuits, and cortical circuits in general, function has been largely based on how fast synaptic glutamatergic and GABAergic transmission shapes inter-cellular communication (Anastasiades and Carter, 2021; Kepecs and Fishell, 2014; Harris and Shepherd, 2015; Naka and Adesnik, 2016). Cortical circuits, including the mPFC, are enriched in neuropeptides and their cognate receptors (Smith et al., 2019; Brockway and Crowley, 2020; Casello et al., 2022; Zhong et al., 2022). The expression of neuropeptides has been important in advancing knowledge of how mPFC circuits are organized and for dissecting cortical circuitry. Specifically, neuropeptides have been exploited as markers of specific cortical cell types and neuropeptide genes have provided experimental means to gain genetic access to sub-populations of cortical neurons. However, there is a paucity in studies that have examined the role of neuropeptides in modulating mPFC circuit function, resulting in a critical knowledge gap in our understanding of the role of neuropeptide transmission in influencing how mPFC circuits integrate information from long-range afferent inputs with local computations carried within mPFC excitatory and inhibitory local-circuit motifs.

Opioid peptides and receptors, including kappa-opioid receptors (KORs) and their endogenous ligand dynorphin (Dyn), are expressed in mPFC circuits (Brockway and Crowley, 2020; Casello *et al*., 2022; Zhong *et al*., 2022; Sohn et al., 2014). Dyn in the mPFC has been implicated in promoting negative affect and anxiety-like behavior, alcohol and morphine dependence-induced cognitive deficits, and stress-induced increases in cardiovascular function (Tejeda et al., 2013; Tejeda et al., 2015; Abraham et al., 2021; Fassini et al., 2015; Wei et al., 2022; Wall and Messier, 2002). Further, KOR phosphorylation (an index of recent KOR activation) is observed in the mPFC after various stressors (Abraham *et al*., 2021). Collectively, these results suggest that Dyn / KOR signaling in mPFC circuits may change the function of the mPFC to influence stress- and affective behaviors. KOR activation inhibits release of dopamine in mPFC as well as glutamatergic EPSPs originating from amygdala, but not hippocampal, inputs to the mPFC (Tejeda *et al*., 2013; Tejeda *et al*., 2015). These observations suggest that Dyn / KOR system recruitment may serve to modify how mPFC circuits incorporate excitatory drive from distinct afferent inputs, but this possibility has not been directly tested. Interestingly, KOR activation inhibits elevations in extracellular GABA driven by glutamate reuptake blockade in the mPFC, without modifying basal GABA levels (Tejeda *et al*., 2013). This raises the possibility that KORs regulate recruitment of inhibitory networks by excitatory synaptic inputs. Thus, Dyn / KOR signaling may likely regulate mPFC function in a more complex manner beyond simply inhibiting afferent inputs to the mPFC. However, it is unclear how the Dyn / KOR system integrates itself into limbic-cortical circuits and impacts excitatory and inhibitory circuit motifs that regulate information flow into and out of the mPFC. Specifically, it is unclear 1) whether the Dyn / KOR system homogenously regulates all afferent inputs to the mPFC, 2) how it may influence afferent-driven recruitment of mPFC inhibitory sub-networks, and 3) how this system shapes the way the mPFC integrates excitatory transmission from afferent inputs with polysynaptic inhibition to gate spike outputs. Here, we address these gaps using a combination of whole-cell slice electrophysiology, viral and genetic approaches, optogenetics, and two-photon calcium imaging. In this study, we reveal that the expression of functional KORs, or lack thereof, in afferent inputs fundamentally changes the impact of Dyn / KOR signaling on input-output transformations in mPFC pyramidal neurons. This study advances our understanding of how neuropeptidergic signaling interfaces with pathway- and cell type-specific excitatory and inhibitory motifs that shape mPFC information processing.

## Results

### Dyn / KOR Pathway-specific regulation of limbic excitatory inputs to the mPFC

The basolateral amygdala (BLA), paraventricular nucleus of the thalamus (PVT), contralateral PFC (clPFC), and ventral hippocampus (VH) are major limbic inputs to the mPFC. To determine whether Dyn signaling via KOR inhibits afferents to the mPFC in a pathway-specific manner, we injected WT mice with AAV-CaMKII-ChR2-eYFP into the basolateral amygdala (BLA), paraventricular nucleus of the thalamus (PVT), contralateral PFC (clPFC), or ventral hippocampus (VH) and recorded optogenetically-evoked excitatory post-synaptic currents (oEPSCs) onto pyramidal neurons (Fig. 1A). Dyn inhibited excitatory outputs in a pathway-specific manner, including excitatory synapses from the PVT, BLA, clPFC, but not the VH (Fig. 1B, C). Dyn failed to modify spontaneous excitatory post-synaptic current (sEPSC) frequency and amplitude onto layer V pyramidal neurons, suggesting that the majority of excitatory synapses innervating layer V pyramidal neurons are not modulated by presynaptic KOR signaling and that KOR activation does not have a post-synaptic effect on glutamate receptor function (Fig. S1A). Also consistent with a presynaptic site of action, Dyn failed to modify excitatory currents evoked by glutamate uncaging (Fig. S1B). We subsequently injected AAVrg-FLEX-tdTomato in the mPFC of KOR-Cre mice to determine whether differential KOR expression may underlie the pathway-specific inhibition of EPSCs that we observed (Fig 1D). Retrogradely-labeled tdTomato cells that project to the mPFC were observed in the BLA, PVT, clPFC and several higher-order telencephalon and midbrain regions, including the insular cortex, clPFC, piriform cortex, claustrum, perirhinal cortex, periamygdaloid cortex, and ventral tegmental area (VTA; Fig 1D, E; Fig S1C). Importantly, the VH lacked tdTomato-labeled neurons, consistent with a lack of Dyn regulation of VH inputs to the mPFC. Further, KOR mRNA expression was absent in the VH formation of WT mice where outputs to the mPFC originate but present in the PVT and BLA, consistent with lack of KOR-positive VH afferents to the mPFC (Fig. S1D).

**Fig. 1:**
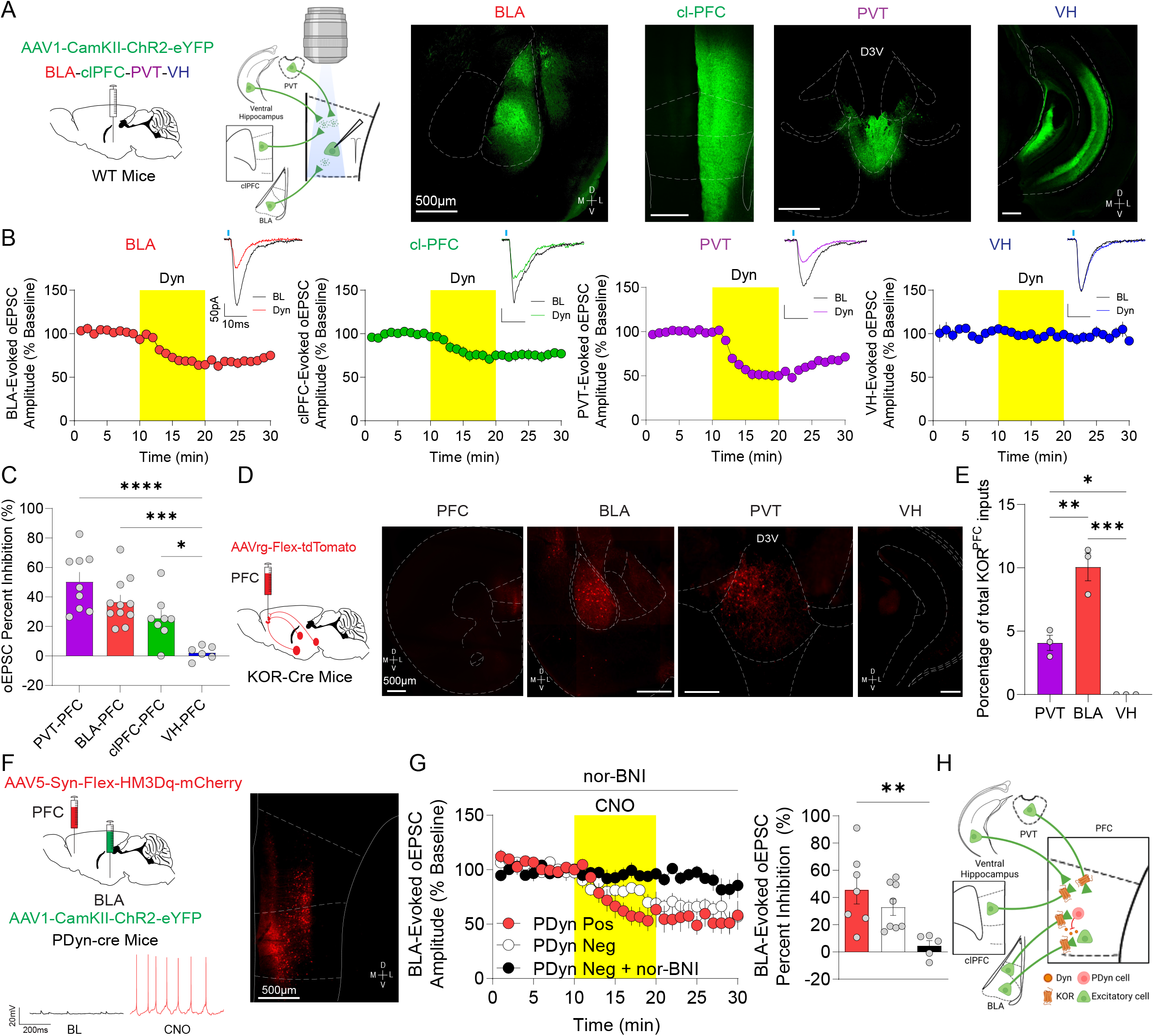
Pathway-specific regulation of mPFC excitatory afferents by the Dyn / KOR system. A. Schematic diagram of the experimental design and representative image AAV1-CaMKII-ChR2-eYFP expression in the PVT, BLA, contralateral PFC, and VH. B. Time course of the effect of KOR activation with Dyn (1 µM) on oEPSC amplitude (expressed as a percentage of baseline) in the mPFC from animals expressing ChR2-eYFP in the PVT (n = 9 cells), BLA (n = 12 cells), clPFC (n = 8 cells) or VH (n = 6 cells). Representative oEPSCs recorded in PFC from PVT (purple traces), BLA (red traces), clPFC (green traces), and VH (blue traces) during baseline (dark traces) and after Dyn (color traces) are shown. Data in this graph and the rest of the manuscript are presented as the mean ± SEM C. Percent inhibition of oEPSC amplitude evoked from different mPFC inputs by Dyn (ANOVA, F_(3, 30)_ = 11.95, p < 0.0001; *p = 0.0471, ***p = 0.0009, ****p < 0.0001). D. KOR-Cre mice were injected unilaterally with AAVrg-FLEX-tdT into the mPFC. Representative images showing retrogradely-labeled tdTomato-expressing KOR-positive cells in different mPFC input areas (clPFC, BLA, VH, and PVT) (ANOVA, F_(2, 6)_ = 49.32, p = 0.0002; *p = 0.0168, **p = 0.0026, ***p = 0.0002). E. Percentage of total KOR-containing projecting cells between different mPFC inputs (n = 3 mice). F. AAV1-CaMKII-ChR2-eYFP expression in the BLA and AAV5-Syn-FLEX-HM3Dq-mCherry in the mPFC of PDyn-Cre mice. Representative traces of CNO-induced activation excitatory DREADD in a mPFC mCherry-positive PDyn cell. G. Time course of the effect of CNO activation (1 µM) on BLA-evoked oEPSC amplitude in PDyn-positive (n = 7 cells of 5 mice) and negative (n = 8 cells) mPFC cells in the presence or absence of nor-BNI (1 µM) (n = 5 cells). Percentage of oEPSC inhibition induced by CNO application and co-application with nor-BNI (ANOVA; F_(2, 17)_ = 6.22, p = 0.0094; **p = 0.0074). H. Pathway-specific KOR modulation of excitatory synapses in the mPFC. KORs inhibit BLA, clPFC, but not VH, afferents onto mPFC neurons.

The source of Dyn acting on presynaptic KOR originating from BLA, PVT, and clPFC is unclear. Indeed, injection of PDyn-Cre mice with AAV expressing Cre-dependent Synaptophysin-GFP-2A-tdTomato revealed that mPFC Dyn neurons have extensive local connections within the mPFC (Fig. S1E). To determine whether endogenous dynorphin release can regulate afferent inputs to the mPFC, we injected AAV-CaMKII-ChR2-eYFP into the BLA and Cre-dependent HM3Dq-expressing virus into the mPFC of PDyn-Cre mice (Fig. 1F). Activation of mPFC Dyn cells using CNO produced robust firing that persisted during drug application. CNO bath application inhibited BLA-evoked oEPSCs onto HM3Dq-positive Dyn neurons or Dyn-negative counterparts (Fig 1G). This effect was not present in cells pretreated with the KOR antagonist nor-BNI. These results suggest that Dyn acts as a negative feedback signal on KOR-positive inputs onto mPFC Dyn cells. Endogenous Dyn inhibition of KOR-positive inputs from the BLA in mPFC Dyn negative cells is consistent with the hypothesis that Dyn acts via volume transmission in mPFC circuits. These results demonstrate that mPFC Dyn cells can broadly influence excitatory afferent drive of mPFC neurons in a pathway-specific manner onto both Dyn positive and negative neurons (Fig. 1H).

### Dyn / KOR signaling distinguishes parallel pathways innervating the mPFC

BLA and mPFC neurons are heterogenous and consists of cell types that differ based multiple features, including molecular identity (Tye and Janak, 2007; Kim et al., 2017; Zhang et al., 2021; Tasic et al. 2018). It is possible that KOR expression and regulation of glutamate release may be an additional defining feature that may differentiate intermingled heterogenous populations of amygdalo-cortical and cortico-cortical neurons. To determine whether functional KOR regulation differentiates parallel ascending amygdalocortical pathways, we injected KOR-Cre mice with virus expressing ChR2 in the BLA in a Cre-ON- or Cre-OFF-dependent manner to express ChR2 in KOR-containing and KOR-lacking BLA neurons, respectively (Fig. 2A). Dyn inhibited optogenetically-evoked excitatory inputs from BLA synapses in Cre-ON ChR2-expressing mice, but not mice expressing Cre-OFF ChR2 (Fig. 2B). This provides conclusive evidence that the effects of Dyn on BLA inputs occurs via the KOR receptor that is localized to presynaptic BLA terminals. Furthermore, KOR-expressing and lacking BLA inputs to the mPFC were sensitive to enkephalin, a distinct opioid peptide with a low affinity for KORs (Fig. S2A, B). These results suggest that parallel amygdalo-cortical pathways are differentiated by dynorphin/KOR signaling, but not enkephalin sensitivity. They also confirm the specificity for dissecting KOR responsivity with Cre-ON and Cre-OFF approaches. We further determined whether KOR regulation may segregate parallel pathways arising from the clPFC (Fig. 2A, C). Dyn inhibited oEPSCs evoked by stimulation of KOR-positive, but not KOR-negative, inputs arising from the clPFC (Fig 2C), suggesting that Dyn / KOR regulation may contribute to parcellation of subnetworks within various afferent inputs to the mPFC. Injection of Cre-OFF ChR2 virus into the PVT did not yield sufficient expression in the mPFC to perform similar experiments to those described above (data not shown), which precluded our ability to examine KOR-positive and KOR-negative PVT inputs to the mPFC using this approach. Retrograde labeling of mPFC-projecting cells utilizing retrobeads demonstrated that mPFC-projecting cells in the BLA, PVT, clPFC expressed or lacked KOR mRNA (Fig 2D, E, F), providing further evidence of parallel pathways that differ based on KOR expression. Consistent with our previous experiments (Fig 1), KOR mRNA was not detected in VH neurons that innervate mPFC. Parallel inputs to the mPFC from KOR Cre mice injected with Cre-ON and Cre-OFF anterograde tracers primarily innervated the ventral mPFC (vmPFC; Fig. 2G, H). However, KOR-positive inputs innervated more dorsal aspects of the vmPFC and dmPFC and deeper mPFC layers relative to KOR-lacking inputs. Additionally, KOR-negative terminals were also observed in the orbitofrontal cortex, which was not observed for KOR-positive neurons. Collectively, these results demonstrate that presynaptic KOR sensitivity reveals the existence of parallel networks innervating the PFC and that Dyn release from sub-populations of mPFC neurons selectively refines excitation from KOR-positive sub-networks embedded within select afferent inputs.

**Fig. 2:**
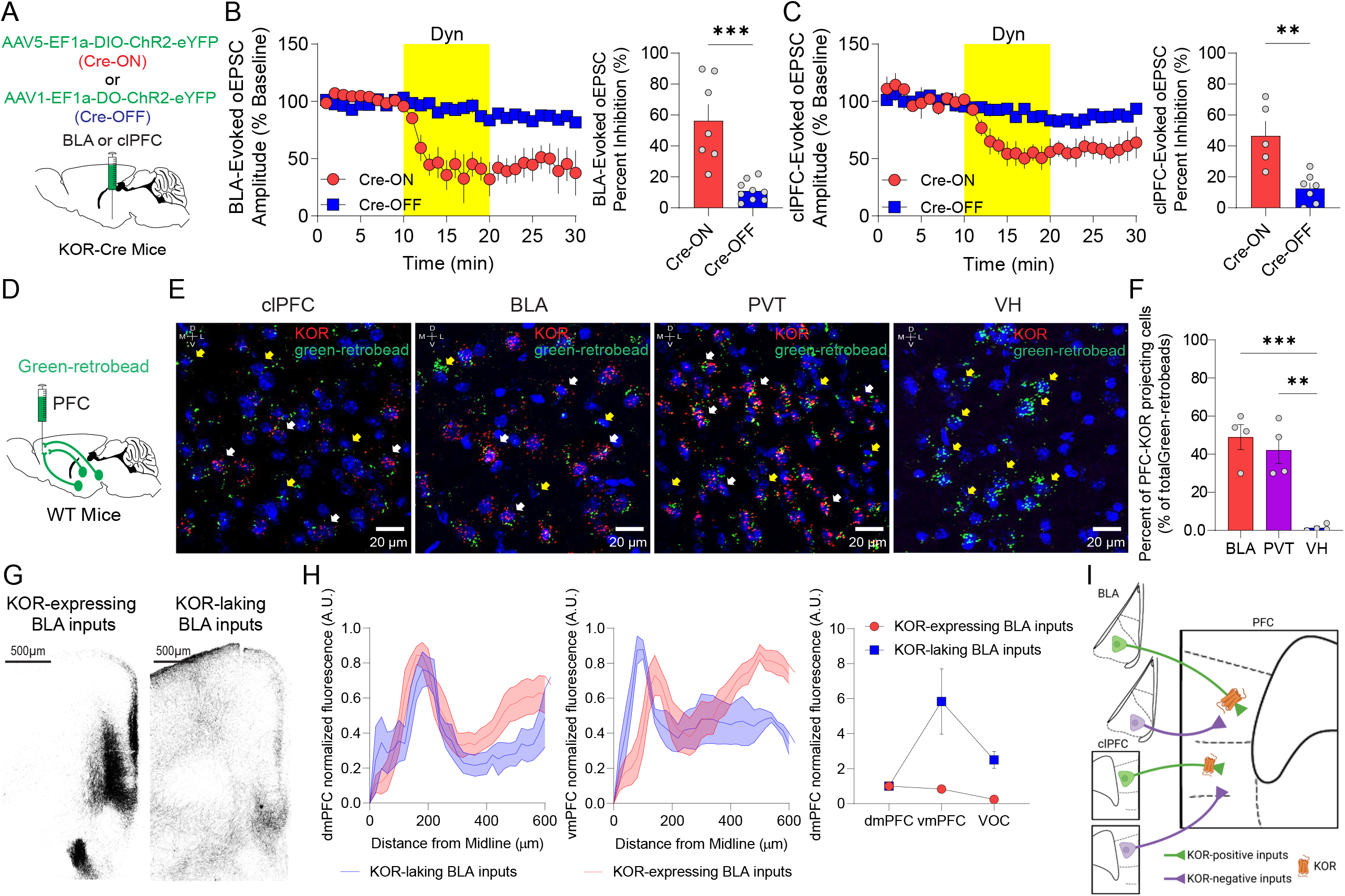
Functional Dyn / KOR signaling distinguishes parallel pathways innervating the mPFC. A. KOR-Cre mice were injected bilaterally with AAV5-EF1α-DIO-ChR2-eYFP (Cre-ON) or AAV1-EF1α-FLOX-ChR2-eYFP (Cre-OFF) into the BLA or contralateral PFC. B. Time course of Dyn (1 µM) regulation of BLA-evoked oEPSC amplitude in the mPFC principal neurons from animals expressing ChR2-eYFP in KOR-containing (Cre-ON) (n = 8 cells) or KOR-lacking (Cre-OFF) (n = 10 cells; Two-way ANOVA; Time x virus interaction; F_(24, 352)_ = 12.36, *p* < 0.0001) neurons in BLA. Percentage of oEPSC inhibition by Dyn application between KOR-containing (Cre-ON) and KOR-lacking (Cre-OFF) ChR2-eYFP in the BLA (t_(14)_= 4.80, p = 0.0003). C. Same as B, but in animals expressing ChR2-eYFP in KOR-positive (Cre-ON) (n = 7 cells) or KOR-negative (Cre-OFF) (n = 10 cells) neurons in the clPFC (Time course Two-way ANOVA; Time x virus interaction; F_(24, 240)_ = 5.029, *p* < 0.0001; Percent inhibition t-test; t_(10)_= 3.88, p = 0.003). D. Wild-type mice were injected bilaterally with green-retro beads into the PFC. E. Representative image of KOR mRNA (red puncta), retro beads (green puncta), and DAPI (blue) in mPFC inputs areas (contralateral PFC, Basolateral Amygdala, paraventricular thalamus, and ventral hippocampus). F. Quantification of colocalization of KOR mRNA and green-retro beads in PVT, BLA, and VH (n = 4 mice, ANOVA, F_(2, 9)_ = 21.21, p = 0.0004; **p = 0.0015, ***p = 0.0005). G. Representative image of KOR-containing and KOR-lacking BLA inputs across mPFC subregions and layers. H. Quantification of KOR-containing (n = 4 mice) and KOR-lacking (n = 5 mice) fibers from the BLA across dmPFC (Two-way ANOVA; Distance x virus interaction; F_(31, 279)_ = 1.463, *p*=0.059) or vmPFC layers (Two-way ANOVA; Distance x virus interaction; F_(31, 279)_ = 5.81, *p*<0.0001). Fluorescence quantification of KOR-positive and KOR-negative BLA inputs in dmPFC, vmPFC, or mOFC (Two-way ANOVA; virus X PFC sub-region interaction, F_(2, 21)_ = 3.94, *p* = 0.0352). I. Parallel pathway-specific KOR modulation of BLA excitatory synapses in the PFC. KOR inhibits BLA and clPFC KOR-expressing inputs to the mPFC, but not BLA and clPFC KOR-negative fibers.

### Dyn / KOR signaling inhibits afferent-driven polysynaptic inhibition in a pathway-independent manner

Afferent inputs to the mPFC recruit polysynaptic inhibition to shape mPFC circuit function (Anastasiades and Carter, 2021). Since Dyn / KOR signaling inhibits the ability of glutamate reuptake inhibition to elevate extracellular GABA levels (Tejeda *et al*., 2013), it is possible that Dyn / KOR signaling may regulate polysynaptic inhibition in the mPFC recruited by afferent inputs. To test this, we injected AAV expressing ChR2 into BLA, VH, PVT or clPFC and recorded biophysically-isolated optogenetically-evoked polysynaptic IPSCs (opsIPSCs) onto layer V pyramidal neurons (Fig. 3A). Unlike the pathway-specific Dyn inhibition of monosynaptic EPSCs described above (Fig 1 and 2), Dyn inhibits polysynaptic inhibition in mPFC pyramidal neurons in a pathway-independent manner (Fig. 3B), including polysynaptic inhibition driven by inputs from the VH, which do not express functional KORs (Fig. 1). Inhibition of opsIPSCs was more robust than inhibition of excitatory afferents (Fig. 3C). A selective KOR agonist similarly inhibited afferent-driven polysynaptic inhibition (Fig. S3A). To test whether Dyn directly impacts GABA receptor function pyramidal neurons, we measured inhibitory currents in evoked by GABA uncaging and found that these currents were not modified by Dyn application (Fig. S3B) which indicates that KOR activation does not modify post-synaptic GABA receptor function. Furthermore, a KOR agonist failed to modify GABAergic miniature IPSC frequency (mIPSCs; Fig. S3C) suggesting that KOR activation does not modify GABA release probability from a sufficiently large set of inhibitory synapses for us to detect the impact simply by measuring mIPSC frequency. In contrast, Dyn decreases spontaneous IPSC (sIPSC) frequency and sIPSC amplitude (Fig. S3D). Collectively, these results suggest that Dyn does not modify GABA transmission via a post-synaptic site of action or through direct actions on a significant population of inhibitory neurons, but rather through an effect of excitation on mPFC networks.

**Fig. 3:**
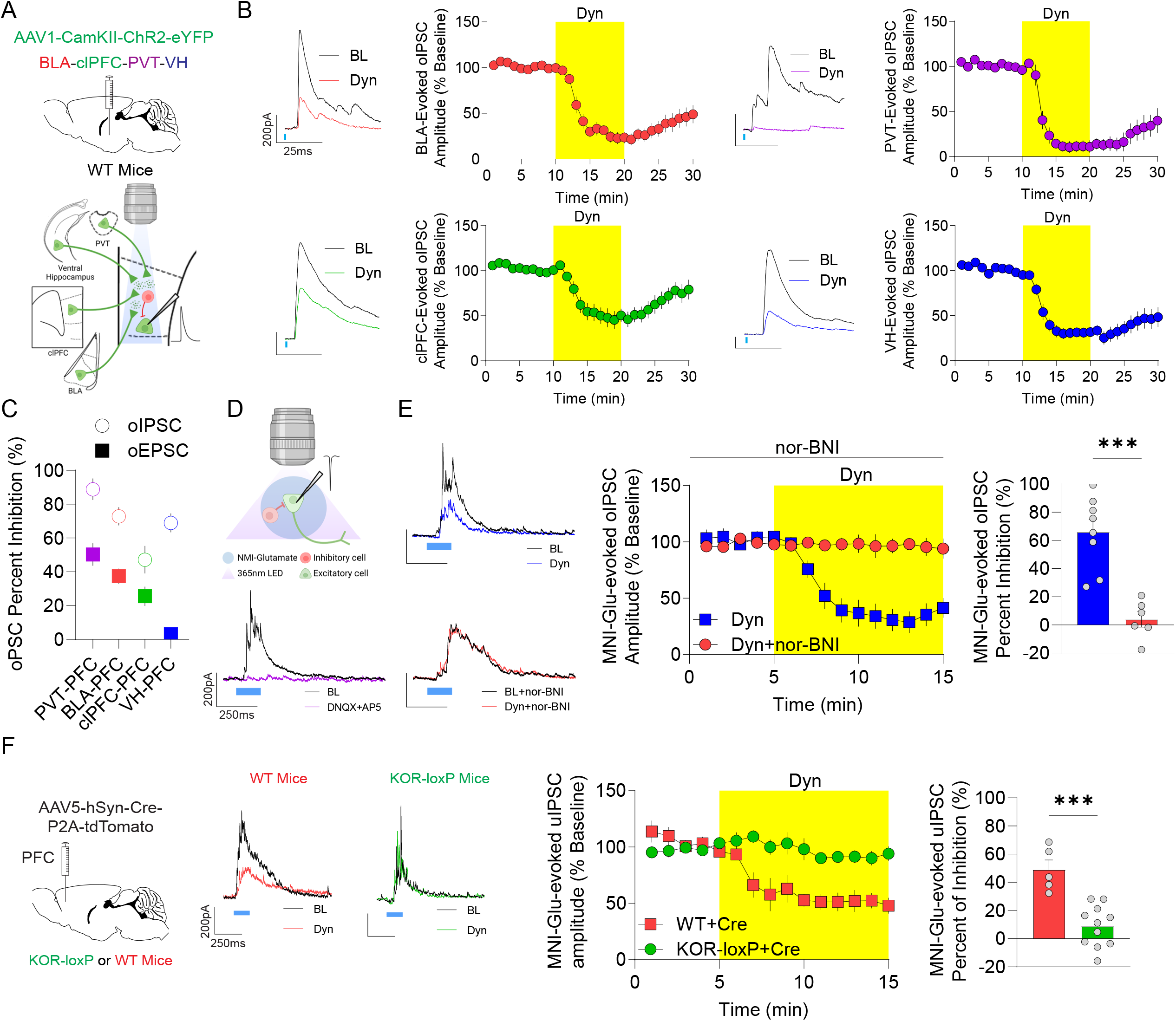
Dyn / KOR signaling inhibits afferent-driven polysynaptic inhibition in a pathway-independent manner. A. Schematic diagram of the experimental design. AAV1-CaMKII-ChR2-eYFP expression in the PVT, BLA, contralateral PFC, and VH. B. Time course of the effect of KOR activation with Dyn (1 µM) on the amplitude of oIPSC in the mPFC from animals expressing ChR2-eYFP in the PVT (n = 9 cells), BLA (n = 9 cells), clPFC (n = 10 cells) or VH (n = 8 cells). Representative oIPSCs recorded in the mPFC evoked by stimulation of PVT (purple traces), BLA (red traces), clPFC (green traces), and VH (blue traces) inputs during baseline (dark traces) and after the KOR agonist, Dyn (color traces). C. Percentage inhibition of polysynaptic oIPSCs and direct oEPSCs by Dyn application between different mPFC inputs (Two-way ANOVA; PSC x pathway interaction; F_(3, 60)_ = 4.37, *p* = 0.0075; PSC main effect; F_(1, 60)_ = 91.94, *p* < 0.0001). D. Schematic diagram and representative oIPSCs were evoked in mPFC principal neurons using MNI-Glutamate. Glutamate uncaging-evoked IPSCs are shown during baseline (dark traces) and after DNQX+AP5 (purple), Dyn (blue), or Dyn+norBNI (red). E. Time course of the effect of KOR activation with Dyn on the amplitude (expressed as a percentage of baseline) of oIPSCs evoked by MNI-glutamate uncaging in the mPFC (n = 8 cells) or co-application norBNI (n = 6 cells; Two-way ANOVA; Time x drug interaction; F_(14, 164)_ = 11.29, *p* < 0.0001). Comparison of percent inhibition of oIPSCs evoked by MNI-glutamate by Dyn and in cells with concomitant Dyn and nor-BNI bath application (t_(12)_= 5.33, *p* = 0.0002). F. KOR-loxP mice were injected bilaterally with AAV5-hSyn-Cre-P2A-tdTomato into the mPFC. Representative trace of oIPSCs evoked by MNI-glutamate uncaging. Traces of IPSCs during baseline (dark traces) and after Dyn in WT (red) or KOR-loxP (green) mice. Time course of the effect of KOR activation on the amplitude of oIPSCs evoked by MNI-glutamate in mPFC neurons from WT (n = 5 cells) or KOR-loxP (n = 11 cells) mice (Two-way ANOVA; Time x virus interaction; F_(14, 194)_ = 8.44, *p* < 0.0001). Percent inhibition of oIPSC induced by MNI-glutamate by Dyn in WT and KOR-loxP mice (t_(14)_= 5.01, p = 0.0002).

### Dyn / KOR signaling within mPFC circuits contributes to Dyn-mediated disinhibition

Strong inhibition of polysynaptic inhibition evoked by VH afferents to the mPFC despite a lack of effect on direct excitatory inputs indicates that Dyn / KOR-mediated disinhibition may be mediated via Dyn / KOR signaling within mPFC local circuits. To test whether Dyn regulates recruitment of polysynaptic inhibition via actions on local circuitry, we uncaged glutamate to bypass excitatory inputs to the mPFC and examined glutamate uncaging-evoked polysynaptic IPSCs, which were completely blocked by the AMPA and NMDA receptor antagonists DNQX and AP5, respectively (Fig 3D). Glutamate uncaging-evoked psIPSCs were robustly inhibited by Dyn, an effect that was blocked by the KOR antagonist nor-BNI (Fig. 3E). To determine whether expression of KORs by mPFC neurons mediates the disinhibitory effects of Dyn, we genetically ablated KORs within mPFC circuits using KOR loxP mice and site-directed Cre expression (Fig. 3F). Dyn failed to inhibit glutamate uncaging-evoked psIPSCs in KOR loxP mice injected with intra-mPFC AAV-Cre, an effect that was intact in Cre-expressing WT controls (Fig. 3F). This demonstrates that Dyn / KOR signaling within mPFC circuits contribute to disinhibition. These results also further demonstrate the specificity of Dyn on the KOR to disinhibit mPFC circuits and provide a functional verification of successful KOR genetic ablation. Collectively, these results suggest that Dyn disinhibits mPFC pyramidal neurons by activating KORs embedded within mPFC circuits.

### mPFC KOR-expressing excitatory neurons engage local circuit inhibition

We further sought to dissect the impact that Dyn / KOR signaling may have on intra-mPFC inhibitory motifs that underlie Dyn-mediated disinhibition (Fig. 3). If Dyn / KOR signaling within mPFC circuits promotes disinhibition, then KOR-expressing neurons are predicted to be embedded within mPFC circuits. One possibility is that Dyn / KOR signaling acts as an inter-cellular communication motif via differential expression of Dyn and KOR across mPFC cell types. An alternative, but not mutually exclusive, possibility is that the Dyn / KOR system functions as an autocrine signal in cells co-expressing Dyn and KOR. Pdyn and KOR mRNA expression did not overlap within mPFC circuits (Fig 4A). Specifically, Pdyn mRNA-positive neurons were localized more superficially than KOR mRNA positive neurons (Fig 4B). These results suggest that Dyn / KOR signaling may act as a scaffold for selective interactions between distinct neurons within a mPFC microcircuit. Further, KOR mRNA was observed in a sparse set of excitatory and inhibitory neurons, but primarily expressed by excitatory neurons of the mPFC, with sparser labeling in VGAT-expressing neurons (Fig 4C, D). KOR-positive excitatory and inhibitory neurons were localized to deeper layers than KOR-negative counterparts in both the dmPFC (prelimbic) and vmPFC (infralimbic; Fig. S4A, B), demonstrating that KOR-containing excitatory neurons represent a sub-population of mPFC principal neurons and interneurons.

**Fig. 4:**
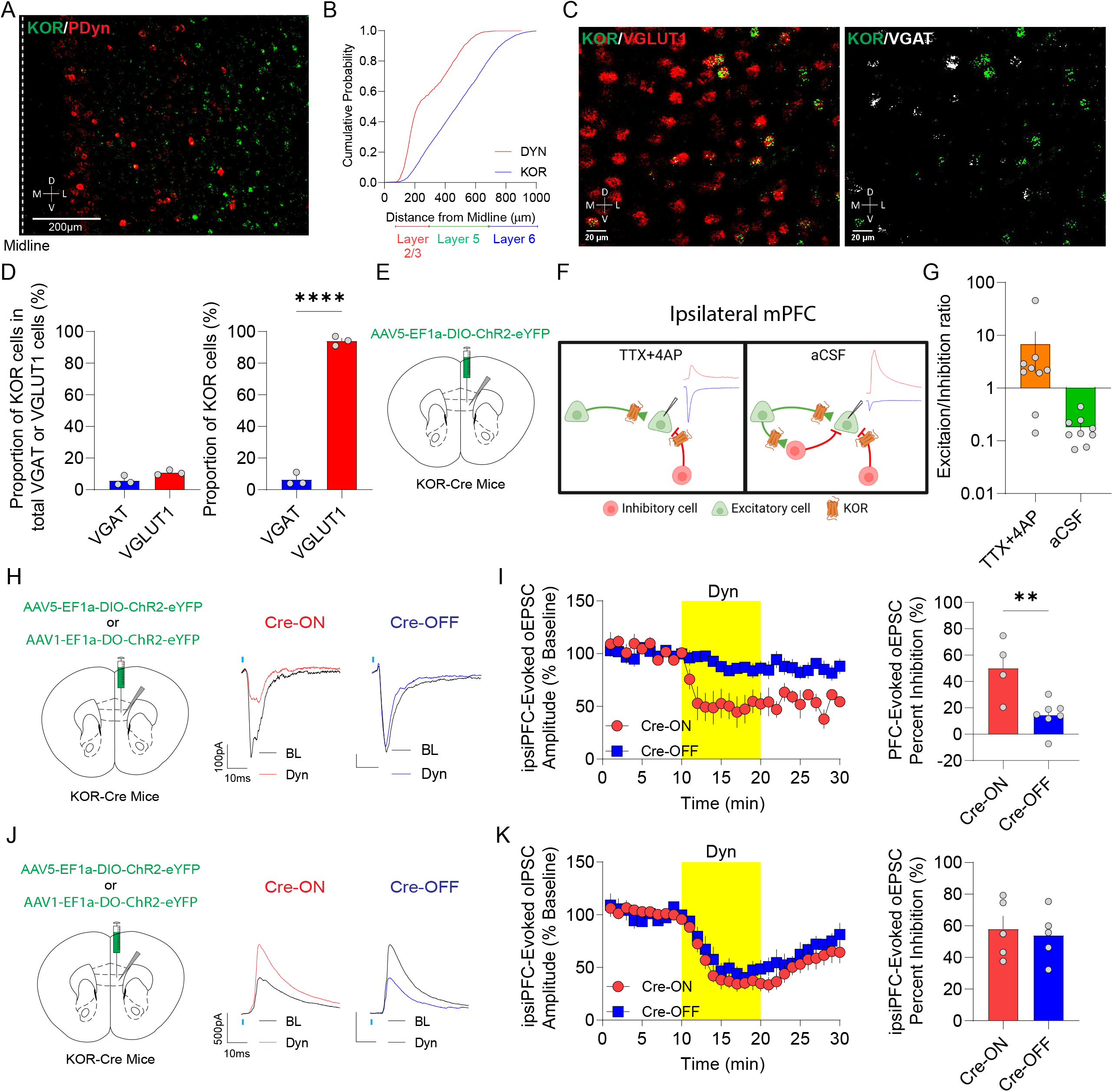
Dyn inhibits mPFC KOR-expressing neuron engagement of feedback inhibition. A. Representative image of KOR mRNA (green puncta) and PDyn mRNA (red puncta) in the mPFC of a WT mouse (n = 4 mice). B. Cumulative probability of KOR and PDyn mRNA across PFC layers (n = 4 mice; K-S Test; D=12.09, *p*<0.0001). C. Representative image of KOR mRNA (green puncta) and VGLUT1 mRNA (red puncta) and VGAT mRNA (white puncta) in the mPFC of a WT mouse (n = 4 mice). D. Mean percentage of KOR mRNA-positive cells expressing in total VGAT and VGLUT1 mRNA (n = 3 mice, t_(4)_= 2.63, *p* = 0.0582). Proportion of total KOR mRNA-positive cells that co-express VGAT and VGLUT1 mRNA (n = 3 mice, t_(4)_= 30.21, *p* < 0.0001). E. KOR-Cre mice were injected unilaterally with AAV5-EF1α-DIO-ChR2-eYFP into the mPFC. F. Schematic of electrophysiological analysis of monosynaptic connectivity by mPFC KOR-positive neurons or polysynaptic recruitment of inhibition. Representative traces of oEPSC (blue) and oIPSC (red) evoked by optogenetic stimulation of mPFC KOR-containing cells. G. Comparison of excitation/inhibition ratios evoked by mPFC KOR cell stimulation in the presence of TTX+4AP (n = 9 cells) or aCSF (n = 9 cells). H. Schematic showing that KOR-Cre mice were injected unilaterally with AAV5-EF1α-DIO-ChR2-eYFP (Cre-ON) or AAV1-EF1α-FLOX-ChR2-eYFP (Cre-OFF) into the mPFC. Representative oEPSCs traces from mPFC neurons of animals injected with Cre-ON or Cre-OFF into the mPFC during baseline (dark traces) and after Dyn (color traces) are shown. I. Time course of the effect of Dyn (1 µM) on oEPSC amplitude of mPFC KOR cell evoked oEPSC from animals expressing in KOR-containing (Cre-ON) (n = 4 cells) or KOR-lacking (Cre-OFF; n = 7 cells) ChR2-eYFP in the ipsilateral mPFC (Two-way ANOVA; Time x virus interaction; F_(24, 211)_ = 5.13, *p* < 0.0001). Percentage of oEPSC inhibition by Dyn application between KOR-containing (Cre-ON) and KOR-lacking (Cre-OFF) ChR2-eYFP in the ipsilateral PFC (t_(9)_= 3.48, *p* = 0.0069). J. KOR-cre mice were injected with AAV5-EF1α-DIO-ChR2-eYFP (Cre-ON) or AAV1-EF1α-FLOX-ChR2-eYFP (Cre-OFF) into the mPFC. Representative oIPSCs traces from mPFC neurons from animals injected with Cre-ON or Cre-OFF ChR2 into the mPFC during baseline (dark traces) and after Dyn (color traces) are shown. K. Time course of the effect of Dyn on the amplitude of oIPSCs in mPFC pyramidal neurons from animals expressing Cre-ON ChR2 in KOR-containing (n = 5 cells) or Cre-OFF ChR2 in KOR-lacking (n = 5 cells) in the ipsilateral mPFC (Two-way ANOVA; Time x virus interaction; F_(24, 240)_ = 0.86, *p* = 0.6436). Percentage of oIPSC inhibition by Dyn application between KOR-containing (Cre-ON) and KOR-lacking (Cre-OFF) ChR2-eYFP in the ipsilateral PFC (t_(8)_= 0.36, p = 0.7252).

Given that Dyn has potent disinhibitory effects via actions on mPFC circuits, the observation that KOR was primarily expressed in excitatory neurons was unexpected. Thus, we hypothesized that KOR-positive excitatory neurons recruit inhibitory mPFC circuits and that Dyn in part decreases feedback inhibition carried by KOR-positive neurons. We observed that KOR-positive mPFC neurons established putative synaptic connections throughout mPFC layers using KOR-Cre mice injected with AAV expressing Cre-dependent Synaptophysin-GFP-2A-tdTomato (Fig. S4C). To identify whether excitatory and inhibitory KOR-positive cells established functional monosynaptic connections within local ipsilateral mPFC (ipsiPFC) circuits we recorded biophysically-isolated excitatory and inhibitory synaptic currents in the presence of TTX and 4-AP in ChR2-negative ipsiPFC neurons evoked by optogenetic stimulation of KOR-positive cells in KOR-Cre mice injected with Cre-dependent ChR2 in the ipsilateral mPFC (Fig. 4E, F; Fig. S4D, E). The majority of neurons received excitatory monosynaptic connections from KOR-positive mPFC neurons, while only half received both direct excitatory and inhibitory connections (Fig. S4F). Of cells that received both excitatory and inhibitory connections, excitatory responses were stronger (Fig. 4G), consistent with preferential expression of KOR mRNA in excitatory neurons. To determine whether mPFC KOR-positive neurons engage local circuit inhibition, we recorded oEPSCs/oIPSCs evoked by optogenetic stimulation of KOR-positive neurons in the ipsilateral mPFC in the absence of TTX/4-AP to permit the recruitment of inhibitory networks within the mPFC. In contrast to monosynaptic responses evoked from KOR-expressing neurons, inhibitory responses dominated over excitatory responses (Fig. 4G), demonstrating that excitatory KOR-expressing neurons potently recruit feedback inhibition within mPFC microcircuits. Collectively, these results demonstrate that the majority of KOR-expressing neurons in the mPFC are deeper layer excitatory neurons that robustly engage local circuit feedback inhibition, providing a potential substrate by which Dyn / KOR signaling disinhibits mPFC circuits.

### Dyn inhibits mPFC KOR-expressing neuron engagement of feedback inhibition

Expression of KOR mRNA in excitatory mPFC cells implies that local circuit Dyn / KOR-mediated disinhibition may in part be a consequence of suppression of action potential generation and/or reduced neurotransmitter release from excitatory mPFC KOR cells that recruit local GABAergic interneurons. Dyn / KOR signaling may shape mPFC circuits via KOR-containing cells either by directly hyperpolarizing KOR-expressing neurons or inhibiting neurotransmitter release. Dyn bath application failed to evoke direct hyperpolarizing currents in KOR-expressing mPFC neurons (Fig. S4G). However, all KOR-expressing cells responded with inhibitory hyperpolarizing currents in response to GABA_B_ receptor activation with baclofen (Fig. S4H). This suggest KORs do not likely couple to GIRKs in mPFC KOR-positive cells, though mPFC KOR-positive cells have the capacity to be directly hyperpolarized by Gi-coupled GPCRs.

It is possible that KOR may inhibit glutamate release from excitatory mPFC KOR-positive cells within ipsilateral mPFC microcircuits, similar to what was observed with clPFC connections (Fig. 1 and 2). To determine whether KOR activation regulates excitatory synapses formed by KOR-expressing neurons within local microcircuits, we injected KOR-Cre mice with Cre-dependent virus expressing ChR2-eYFP (Cre-ON) and recorded oEPSCs in ChR2-negative neurons in the ipsilateral mPFC (Fig. 4H). Dyn bath application inhibited evoked excitatory currents in ChR2-negative mPFC neurons via a presynaptic site of action (Fig. 4I). This effect was still present when monosynaptic currents were isolated using TTX/4-AP (Fig. S4I). Moreover, oEPSCs in pyramidal neurons from KOR-Cre mice expressing Cre-OFF ChR2-eYFP were unaffected by Dyn. These results suggest that mPFC KORs inhibit local circuit excitatory connections established by KOR-expressing pyramidal neurons but do not directly hyperpolarize pyramidal neurons. Moreover, KOR-positive mPFC neurons integrate themselves in local circuits and are distinguished from KOR-lacking cells that also integrate within local circuits (Fig. 4F). Feedback inhibition driven by local circuit KOR-expressing and KOR-lacking neurons was potently inhibited by Dyn (Fig 4J, K). These results suggest that Dyn / KOR signaling inhibits excitatory drive of inhibitory microcircuits by KOR-positive cells and that KOR-negative cells engage KOR-sensitive mPFC neurons that contribute to feedback inhibition.

### Dyn / KOR signaling directly inhibits GABAergic transmission from sub-populations of mPFC interneurons

Since a subset of KOR-expressing neurons are inhibitory interneurons, we determined whether KOR mRNA was differentially expressed in PV and SOM neurons. KOR mRNA expression was observed in sub-populations of PV and SOM mRNA-positive neurons (Fig. 5A-C). Further, PV and SOM-immunoreactive neurons were observed in KOR-Cre mice injected with Cre-dependent virus expressing tdTomato (Fig. S5A), and these cells had similar distribution across mPFC layers as KOR mRNA positive cells (Fig. S5B). The relative abundance of KOR mRNA was similar in PV- and SOM-positive neurons, as well as putative excitatory KOR-positive neurons (Fig. S5C). Further, injecting KOR Cre mice with Cre-dependent GFP virus under the control of the mDlx promoter labeled mPFC interneurons with both fast-spiking and non-fast spiking electrophysiological properties, consistent with expression of KOR in a subset of mPFC interneurons (Fig. S5F, G). In contrast, labeling all KOR cells, and hence biasing towards excitatory mPFC KOR cells, in KOR-Cre mice with Cre-dependent tdTomato resulted in labeled cells with homogenous regular spiking properties typical of subsets of excitatory principal neurons (Fig. S5F, G). Putative excitatory KOR-positive cells which had different physiological properties than KOR-positive mPFC interneurons (Fig. S5H). These results are consistent with KOR expression being limited to a sub-population of excitatory and inhibitory cells. Since KOR mRNA is expressed in a sub-population of SOM and PV neurons in the mPFC, Dyn may inhibit the release of GABA from these populations of cells. To explore whether KORs inhibit GABA release from SOM and PV interneurons, we injected SOM-Cre and PV-Cre mice with AAV-DIO-ChR2-eYFP into the mPFC and recorded optogenetically-evoked IPSCs (oIPSCs) in principal neurons (Fig. 5D, F). Dyn inhibited monosynaptic oIPSCs from SOM- and PV-expressing neurons via a presynaptic site of action, an effect blocked by pretreatment with the KOR antagonist nor-BNI (Fig. 5E; Fig S5I, J). The inhibitory effect of Dyn on SST-evoked oIPSCs was significantly larger than the effect on PV-evoked oIPSCs (Fig. 5F), consistent with KOR mRNA expression in a larger sub-population of SST interneurons than PV interneurons. Thus, KORs may disinhibit mPFC circuits through suppression of both SST and PV interneuron-mediated inhibition, with a bias at suppressing SST-mediated inhibition more robustly.

**Fig. 5:**
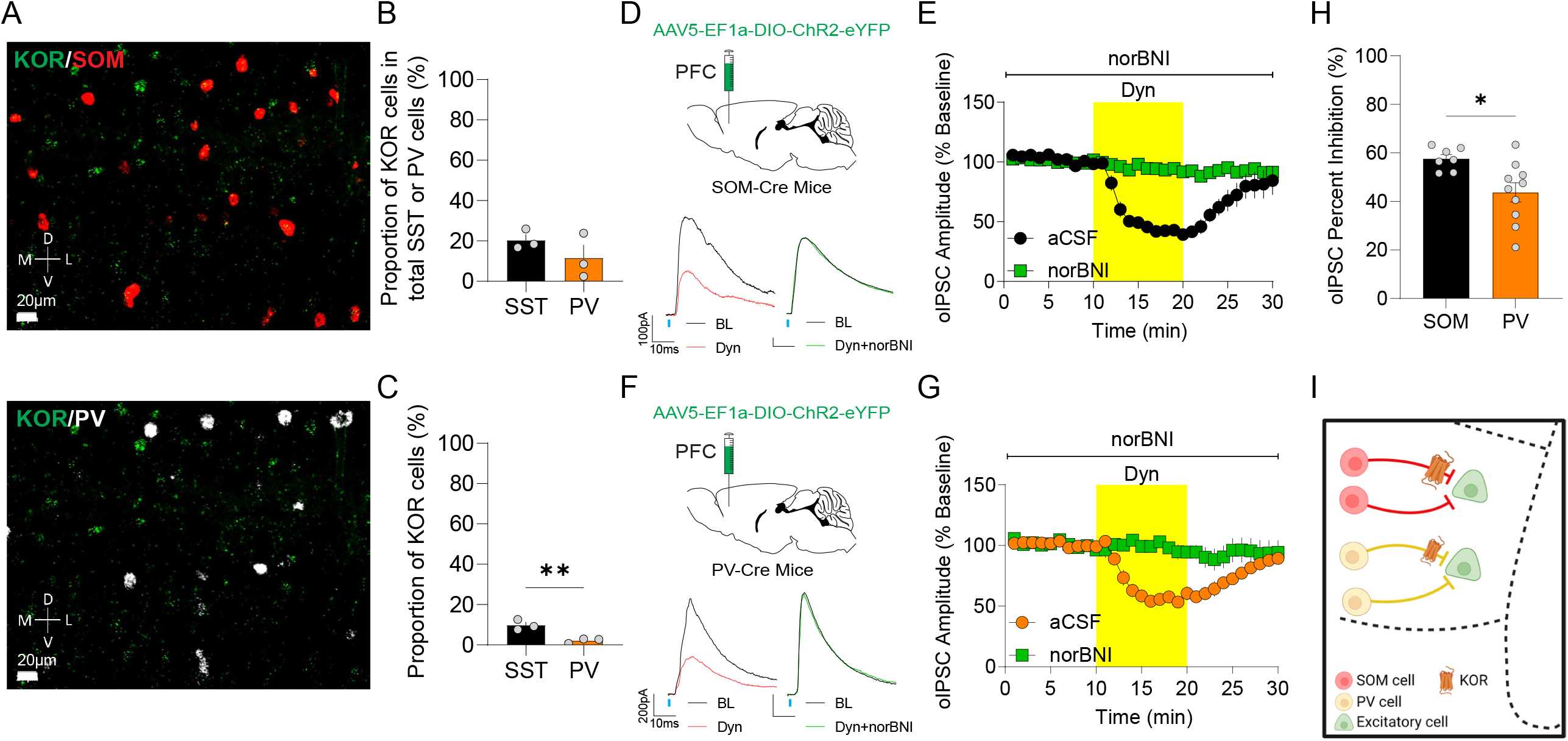
Dyn / KOR signaling directly inhibits GABAergic transmission from sub-populations of mPFC interneurons. A. Representative image of KOR mRNA (green puncta), SOM mRNA (red puncta), and PV mRNA (white puncta) in the mPFC of a WT mouse. B. Mean percentage of KOR mRNA-positive cells expressing in total SOM and PV mRNA (n = 3 mice, t_(4)_= 1.25, *p* = 0.2778). C. Proportion of total KOR mRNA-positive cells that co-express SOM and PV mRNA (n = 3 mice, t_(4)_= 4.69, *p* = 0.0093). D. SOM-Cre mice were injected bilaterally with AAV5-EF1α-DIO-ChR2-eYFP PFC. Traces are shown during baseline (dark traces) and after Dyn (1 μM; red), and Dyn + norBNI (green) are shown. E. Time course of the effect of Dyn on the amplitude of oIPSCs in principal neurons from animals expressing ChR2 in SOM-positive interneurons (n = 7 cells) in the presence or absence of nor-BNI (1 µM) (n = 3 cells; Two-way ANOVA; Time x virus interaction; F_(24, 192)_ = 12.10, *p* <0.0001). F. Schematic depicting the injection of AAV5-EF1α-DIO-ChR2-eYFP into the mPFC of PV-Cre mice. Traces during baseline (dark traces) and after Dyn (red), and Dyn+norBNI (green) are shown. G. Dyn inhibits the amplitude of PV-evoked oIPSC on mPFC pyramidal neurons in the presence (n =10 cells) or absence of nor-BNI (n = 6 cells; Two-way ANOVA; Time x virus interaction; F_(24, 336)_ = 11.55, *p* <0.0001). H. Percentage of oIPSC inhibition by Dyn between PV or SOM evoked by ChR2-eYFP in the mPFC (t_(15)_= 2.75, *p* = 0.0147). I. Model representing that KOR inhibits oIPSCs evoked from SOM- and PV-positive interneurons in the mPFC.

### Dyn / KOR signaling inhibits feedforward inhibition mediated by SST, but not PV, mPFC interneurons

In addition to contributing to disinhibition via KORs expressed within mPFC circuits, presynaptic KORs on afferent inputs may also regulate afferent drive of mPFC interneurons to influence feedforward inhibition. A direct comparison of the effects of Dyn on KOR-positive (e.g. BLA) and negative (e.g. VH) afferents was precluded given that KOR-positive afferent inputs to, including those from the BLA and clPFC, consists of intermingled KOR-sensitive and insensitive fibers. To directly examine whether presynaptic KOR regulation of excitatory afferents contributes to Dyn-mediated disinhibition, we injected KOR-Cre mice with AAV expressing either Cre-ON ChR2 or Cre-OFF ChR2 into the BLA to evaluate polysynaptic oIPSCs evoked by stimulation of KOR-expressing and KOR-lacking BLA inputs in mPFC, respectively (Fig. 6A). In stark contrast to Dyn sensitive- and -insensitive monosynaptic EPSCs arising from KOR-positive (Cre ON) and KOR-negative (Cre-OFF) BLA inputs (see Fig. 2), Dyn inhibition of opsIPSCs was observed in both Cre-ON and Cre-OFF-ChR2-expressing mice (Fig. 6B, C), an effect that was significantly stronger in the KOR-containing BLA input than their KOR-lacking counterparts (Fig. 5B,C). These results suggest that the presence of a presynaptic KOR contributes to Dyn-mediated disinhibition. If this is the case, the result should generalize to other pathways with presynaptic KORs. We next tested this by injecting AAV expressing either Cre-ON ChR2 or Cre-OFF ChR2 into the clPFC (Fig. 6D). opsIPSCs driven by KOR-positive and KOR-negative inputs from the clPFC were both inhibited by Dyn (Fig. 6E, F). However, this effect was more robust in the KOR-positive clPFC afferent inputs than KOR-negative clPFC inputs (Fig. 6E, F). These results suggest that in addition to the disinhibitory effect of KORs expressed by mPFC neurons (Fig. 3), presynaptic KORs reduce synaptic excitation of inhibitory neurons within mPFC and serve as a separate mechanism of Dyn-mediated disinhibition.

**Fig. 6:**
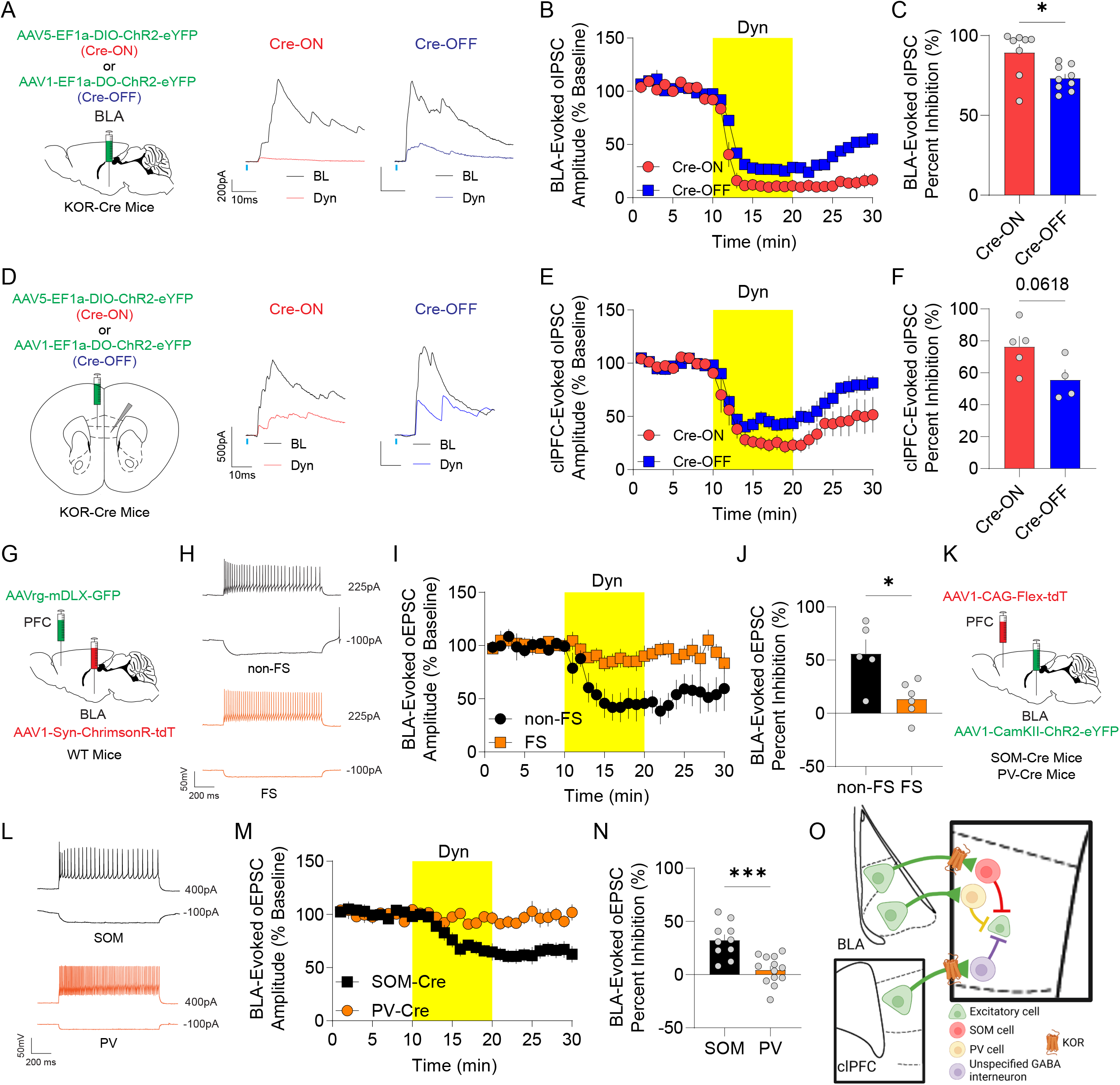
Dyn / KOR signaling inhibits feedforward inhibition by sub-networks of mPFC interneurons. A. KOR-Cre mice were injected bilaterally with AAV5-EF1α-DIO-ChR2-eYFP (Cre-ON) or AAV1-EF1α-FLOX-ChR2-eYFP (Cre-OFF) into the BLA. Representative traces of oIPSCs in mPFC neurons from animals injected with Cre-ON or Cre-OFF ChR2-eYFP into the BLA during baseline (dark traces) and after Dyn (color traces). B. Time course of oIPSC amplitude in mPFC neurons from animals expressing ChR2 in BLA KOR-containing (Cre-ON) (n =8 cells) or KOR-lacking (Cre-OFF) (n = 9 cells) cells (Two-way ANOVA; Time x virus interaction; F_(24, 360)_ = 3.24, *p* <0.0001). C. Percent inhibition of BLA-evoked polysynaptic oIPSCs by Dyn application from KOR-containing (Cre-ON) and KOR-lacking (Cre-OFF) neurons (t_(15)_= 2.86, *p* = 0.0117). D. Schematic depicting injection of KOR-Cre mice with AAV5-EF1α-DIO-ChR2-eYFP (Cre-ON) or AAV1-EF1α-FLOX-ChR2-eYFP (Cre-OFF) into the clPFC and representative traces of oIPSCs during baseline (dark traces) and after the KOR agonist, Dyn1-17 (color traces). E. Same as A but from animals expressing ChR2-eYFP in KOR-containing (Cre-ON) (n = 5 cells) or KOR-lacking (Cre-OFF) (n = 4 cells) in the clPFC (Two-way ANOVA; Time x virus interaction; F_(24, 168)_ = 1.67, *p* = 0.0315). F. Percentage of oIPSC inhibition by Dyn application between KOR-containing (Cre-ON) and KOR-lacking (Cre-OFF) ChR2-eYFP in the contralateral PFC (t_(7)_= 2.22, *p* = 0.0618). G. WT mice were injected bilaterally with AAV1-CaMKII-ChR2-eYFP into the BLA and AAVrg-mDLX-GFP into the PFC. H. Representative traces at different current steps of evoked action potentials in mPFC fast-spiking interneurons (FSIs; orange) and non-fast spiking interneurons (non-FSI; black). I. Time course of the effect of Dyn on the amplitude of BLA-evoked oEPSCs in FSIs (n = 6 cells) or non-fast spiking interneurons (non-FSI) (n = 5 cells; Two-way ANOVA; Time x virus interaction; F_(24, 212)_ = 3.80, *p* <0.0001). J. Dyn-induced inhibition of BLA-evoked oEPSCs (percent inhibition) in mPFC FSIs and non-FSIs (t_(9)_= 3.02, *p* = 0.0143). K. Schematic depicting injection of AAV1-CaMKII-ChR2-eYFP into the BLA and AAVrg-Flex-tdT into the mPFC of SOM-Cre or PV-Cre mice. L. Representative non fast-spiking and fast-spiking properties from SOM-positive (black) and PV-positive (orange) mPFC interneurons, respectively. M. Time course of the effect of Dyn on BLA-evoked oEPSC amplitude in mPFC SOM (n = 11 cells) or PV (n = 13 cells; Two-way ANOVA; Time x virus interaction; F_(24, 504)_ = 6.54, *p* <0.0001). N. Percentage of oEPSC inhibition by Dyn between PV or SOM evoked by ChR2-eYFP positive terminals in the mPFC (t_(21)_= 4.52, *p* = 0.0002). O. Parallel pathway-specific KOR modulation of BLA or clPFC evokes feed-forward inhibition in the mPFC. KOR inhibit BLA inputs onto mPFC non-FSI SOM-positive, but not PV-positive FSIs.

Excitatory drive of diverse GABAergic interneurons by limbic afferent inputs recruits rapid feedforward inhibition that is essential for appropriate mPFC function and control of behavior (Kepecs and Fishell, 2014; Anastasiades and Carter, 2021). To determine whether presynaptic KORs regulate excitatory inputs to distinct GABAergic neurons, we injected WT mice with AAV-hSyn-Chrimson-tdTomato into the BLA and AAV-mDlX-eGFP into the mPFC to label and record from the two largest groups of cortical interneurons, PV- and SOM-positive GABAergic interneurons (Fig. 6G; Dimidschstein et al., 2016). Intrinsic firing properties of GFP-expressing neurons were determined, and cells were divided into fast-spiking and non-fast-spiking neurons (Fig. 6H). Interestingly, Dyn inhibited BLA oEPSCs onto non-fast-spiking mPFC Dlx-positive interneurons, but not Dlx-positive fast-spiking interneurons (Fig. 6I, J). These results suggest that BLA inputs innervating different inhibitory neuron populations within the mPFC may differentially express KORs. Given that non-fast-spiking interneurons may correspond to a plethora of inhibitory neurons, including SOM interneurons, we further explored the role of Dyn in regulating inputs from the BLA onto SOM and PV interneurons, which largely encompass various non-fast spiking and fast-spiking interneuron classes, respectively (Kepecs and Fishell, 2014; Harris and Shepherd, 2015; Kupferscmhidt et al 2022; Anastasiades and Carter, 2021; Fig. 6L). To this end, we injected SOM-Cre and PV-Cre mice with AAV expressing ChR2 into the BLA and AAV-FLEX-tdTomato into the mPFC of mice to specifically record oEPSCs onto SOM and PV neurons, respectively (Fig. 6K). Dyn inhibited excitatory inputs from the BLA onto SOM-expressing, but not PV, interneurons (Fig. 6M, N). Thus, in addition to inhibiting direct excitatory inputs to mPFC principal neurons from KOR expressing inputs, the Dyn / KOR system may also shape feedforward inhibition coming into the mPFC, by inhibiting excitatory synapses onto SOM-positive interneurons, but not PV interneurons (Fig. 6O). Taken together, these results demonstrate that Dyn / KOR signaling disinhibits mPFC circuits via biased regulation of inhibitory subnetworks. Specifically, presynaptic KORs inhibit excitatory drive of SST interneurons and suppress GABA release from subsets of inhibitory KOR-positive SST interneurons, reducing SST-mediated feedforward inhibition via two independent mechanisms. In contrast, there is less Dyn / KOR regulation of PV interneuron-mediated feedforward inhibition as excitatory synapses innervating PV interneurons are not regulated by KORs and KOR mRNA-expressing PV interneurons and KOR-mediated suppression of GABA release is more limited in PV interneurons than SST interneurons.

### Dyn / KOR signaling bidirectionally shapes input-output transformations in a pathway-dependent manner

Here we have discovered that Dyn / KOR signaling decreases excitatory drive of mPFC neurons in a synapse-specific manner via selective KOR expression (between distinct afferent inputs and parallel channels within pathways), but broadly disinhibits mPFC circuits irrespective of whether the afferent input expresses KOR. This raises the possibility that Dyn / KOR signaling produces opposing effects on integration of excitatory and inhibitory synaptic inputs onto mPFC principal neurons when mPFC circuits are being controlled by either KOR-positive and KOR-negative excitatory afferents. To determine whether Dyn / KOR signaling regulated KOR-negative and positive afferent driven recruitment of mPFC ensembles, we used two-photon imaging of mPFC calcium activity in acute mPFC brain slices in response to optogenetic stimulation of VH or BLA inputs, respectively (Fig. 7A). Dyn enhanced VH-driven calcium activity in mPFC networks in mice expressing Chrimson in the VH in comparison to slices treated with vehicle (Fig. 7C). Conversely, Dyn decreased BLA-evoked GCaMP activity relative to vehicle-treated slices (Fig. 7B), consistent with presynaptic inhibition of KOR-positive BLA glutamatergic inputs to mPFC neurons that limits recruitment of mPFC neurons.

**Fig. 7:**
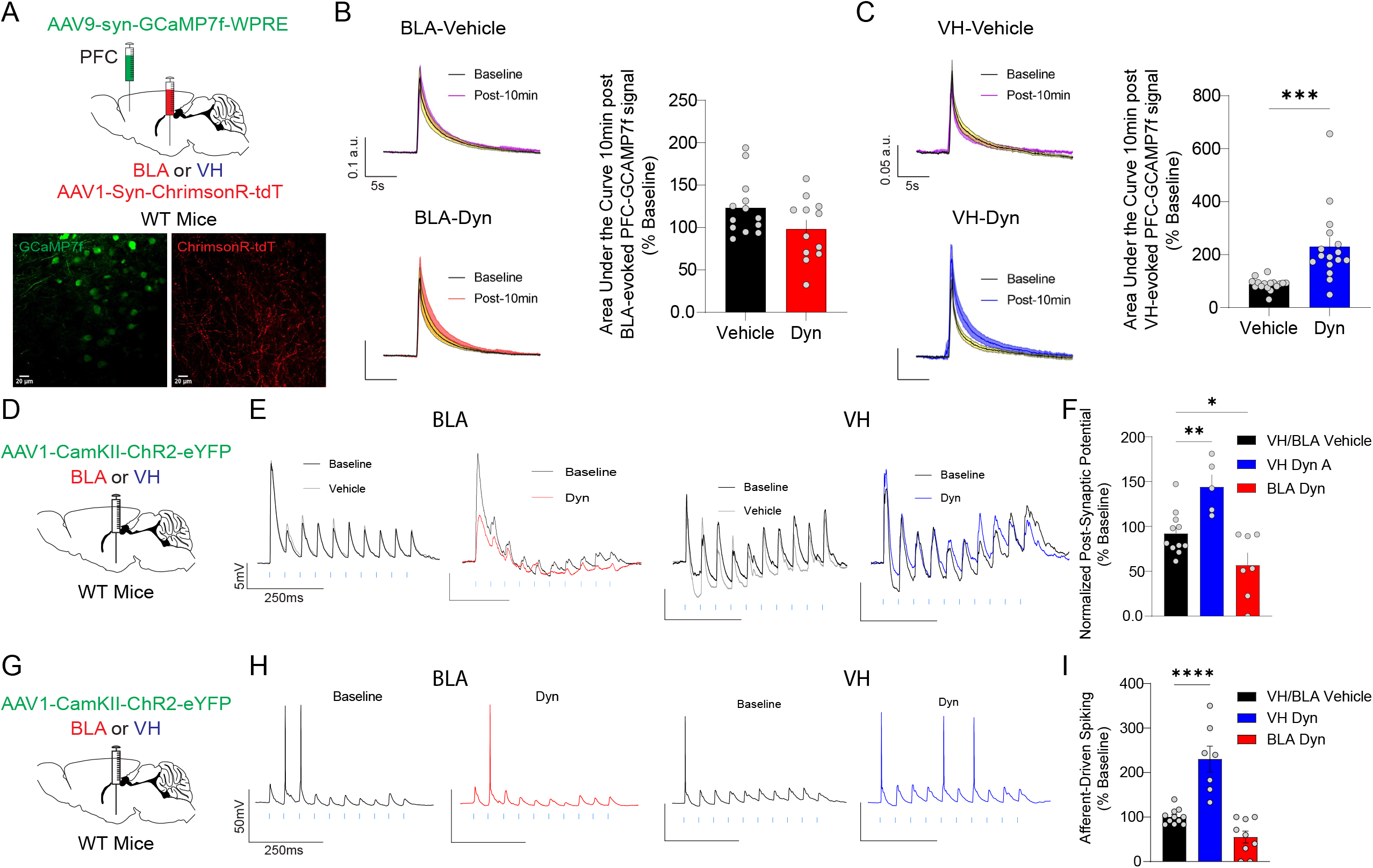
Dyn / KOR signaling bidirectionally shapes input-output transformations in a pathway-dependent manner. A. WT mice were injected bilaterally with AAV1-Syn-ChrimsonR-tdT in the BLA or VH and AAV9-syn-GCaMP7f-WPRE in the PFC. Representative two-photon images of PFC GCaMP7f (green) and ChrimsonR (red) expression in PFC. B. Left, representative traces of GCaMP7f signal for vehicle and Dyn bath application. Right, area under the curve of GCaMP7f signal evoked by a train stimulation of BLA inputs in mPFC slices (n = 13 and n = 12 slice for vehicle and Dyn respectively, t_(23)_= 1.76, *p* = 0.0903). C. Same as for B, but for VH inputs (n = 16 and n = 16 slice for vehicle and Dyn respectively, t_(30)_= 4.00, *p* = 0.0004). D. Experimental scheme wherein WT mice were injected bilaterally with AAV1-CaMKII-ChR2-eYFP in the BLA or VH. E. Representative traces of PSPs in mPFC neurons in response to train stimulation of BLA inputs (10 pulses at 20 Hz), in the presence of vehicle (gray) or Dyn (1 µM; BLA red, n= 19 cells; vehicle = 11 cells, Dyn = 8 cells); VH blue, n= 14 cells (vehicle = 5 cells, Dyn = 9 cells). Blue lines indicate the timing of LED pulses. F. The effect of Dyn / KOR signaling on the area under the curve of PSPs evoked by train stimulation of VH or BLA inputs normalized to baseline (ANOVA, F_(2, 20)_ = 12.78, *p* = 0.0003; *p = 0.0417, **p = 0.0076). G. Schematic depicting that WT mice were injected bilaterally with AAV1-CaMKII-ChR2-eYFP in the BLA or VH. H. Afferent-driving spikes mPFC neurons in response to train stimulation (10 pulses at 20 Hz) of BLA (n = 18 cells; vehicle = 7 cells, Dyn = 11 cells) or VH inputs (n = 12 cells; vehicle = 4 cells, Dyn = 8 cells) during baseline (black traces) and in the presence of Dyn (red or blue traces. Blue lines indicate the timing of LED pulses. I. Mean number of spikes evoked by a train stimulation of VH or BLA inputs in PFC in response to Dyn application as a percentage of baseline firing (ANOVA, F_(2, 24)_ = 29.57, p < 0.0001; ****p < 0.0001).

Subsequently, we performed current clamp recordings in mPFC pyramidal neurons and determined the effects of Dyn bath application on summation of BLA and VH-evoked excitatory post-synaptic potentials (oPSPs) to probe how Dyn influences integration of excitatory input from KOR-positive and negative sources, respectively with local circuit inhibition. Dyn increased integration of VH-evoked oPSPs (area under the curve) via inhibition of hyperpolarizing responses associated with train stimulation (Fig. 7D, E, F). In contrast, Dyn inhibited summation of BLA-evoked oPSPs, by decreasing direct excitation of pyramidal neurons (Fig. 7D, E, F). Given that VH inputs to mPFC neurons are not directly modulated by KORs (Fig 1. EPSCs under voltage clamp), amplification of VH-evoked PSPs suggest that Dyn-mediated disinhibition may be suppressing inhibition that limits temporal summation of EPSPs necessary to trigger action potentials. To determine how Dyn / KOR signaling shapes input / output transformations in mPFC circuits, we performed current clamp recordings in pyramidal neurons and determined the effects of Dyn bath application on evoked spiking driven by optogenetic stimulation of BLA and VH inputs. Evoked spiking was driven by excitatory synapses but gated by polysynaptic inhibition as evoked spiking was abolished and enhanced by the glutamate and GABA-A receptor blockers DNQX/AP-5 and picrotoxin, respectively (Fig. S7). BLA driven spiking was decreased in mPFC neurons by Dyn, with subsets of mPFC neurons showing a decrease in evoked spiking or no change (Fig. 7G, H, I)). Conversely, Dyn significantly enhanced VH-driven spiking (Fig. 7G, H, I)). Since Dyn did not modify VH-evoked EPSCs in voltage clamp experiments (Fig 1), these results suggest that Dyn/KOR suppression of afferent-driven inhibition amplifies integration of excitatory inputs from KOR-negative afferent inputs. In conclusion, Dyn / KOR signaling shapes input / output transformations by limiting mPFC ensemble recruitment by KOR-sensitive afferents and enhancing incorporation of mPFC ensembles by KOR-insensitive afferents, including the BLA and VH, respectively.

## Discussion

Here we demonstrate that the Dyn / KOR system selectively filters incoming excitatory inputs to the mPFC in a pathway-specific manner via presynaptic KORs. Endogenously released Dyn can act via volume transmission to impact KOR-sensitive inputs onto both Dyn-releasing and Dyn-lacking mPFC neurons. Further, we demonstrate that functional physiological effects of Dyn / KOR signaling define sub-networks embedded within amygdalo-cortical and cortico-cortical pathways, providing a mechanism to regulate specific information flow within limbic inputs to the mPFC. Moreover, we demonstrate that mPFC Dyn / KOR signaling robustly disinhibits mPFC circuits via distinct and partially redundant mechanisms. 1) Selective inhibition of excitatory inputs to SOM-positive, but not PV-positive, interneurons. 2) Suppression of feedback inhibition controlled by excitatory mPFC KOR cells. 3) Inhibition of GABA release from subsets of KOR-positive SOM and PV interneurons. Finally, we demonstrate that Dyn / KOR signaling shapes mPFC input-output transformations in a pathway-specific manner by filtering incoming excitatory BLA afferents that are subject to presynaptic inhibition of glutamate release, while amplifying the ability of KOR-insensitive VH inputs to evoke mPFC spiking via KOR-mediated disinhibitory mechanisms. Collectively, our study provides a circuit-based model wherein enhanced Dyn / KOR signaling reconfigures the state of mPFC networks and fundamentally changes how KOR-expressing and lacking afferent inputs control pyramidal neurons.

Dyn / KOR signaling inhibits inputs to the mPFC in a pathway-specific manner, including inputs from the BLA, clPFC, and PVT. Our results demonstrate that VH afferents to the mPFC are KOR-insensitive and this derives from the lack of KOR mRNA expression in VH output regions. This is consistent with previous work demonstrating that systemic KOR activation inhibits BLA-evoked inputs in anaesthetized rats, but not synaptic responses evoked by electrical stimulation of the fimbria / fornix, which carries hippocampal fibers that project to various regions, including the mPFC (Tejeda *et al*., 2015). Dyn fails to significantly inhibit sEPSCs onto layer V pyramidal neurons in mice (present study), consistent with marginal effects of KOR activation of mEPSPs in rat mPFC slices (Tejeda *et al*., 2013). Together, these results suggest that the majority of excitatory inputs onto mPFC neurons are KOR-insensitive. Indeed, mapping of KOR-positive inputs to the mPFC with KOR-Cre mice failed to label neurons in many inputs with projections to the mPFC, including the VH formation, which is devoid of KOR mRNA expression. Selective regulation of KOR-expressing inputs provides Dyn neurons control over incoming information from specific fibers owing to synapse-specific KOR regulation. Further, here we report that endogenous Dyn release acts as a paracrine / retrograde signal that limits incoming excitation from the BLA, and presumably other KOR-expressing afferents. Endogenous Dyn release not only impacts inputs onto Dyn-positive neurons, but also neighboring Dyn-lacking neurons, providing evidence that Dyn may broadly impact mPFC networks via volume transmission. Our unpublished work has demonstrated that mPFC Dyn neurons are activated in response to threats and threat-predictive cues which produces local release of Dyn. Thus, Dyn released during motivationally-charged behaviors may act as a negative feedback mechanism that serves to limit excitatory drive of Dyn-expressing cells, as well as Dyn negative pyramidal neurons.

Presynaptic KOR regulation is a feature that defines sub-populations of excitatory inputs in amygdalocortical and cortico-cortical networks. This provides mPFC Dyn cells the capacity to selectively filter incoming inputs (if they express KORs) in a cell-specific manner even within the same pathway. This may be of relevance to the processing of specific information conveyed to the mPFC from incoming afferents. For instance, in the BLA, valence, actions/outcomes, internal states, and even different modalities within the same valence are encoded by different subsets of cells (Janak and Tye, 2015; Tye and Janak, 2007; Kim *et al*., 2017). KOR-negative and KOR-positive BLA outputs both innervate the mPFC but have differential dorsal/ventral bias within the mPFC, respectively, as well as differential distribution across layers. Moreover, KOR-expressing neurons did not heavily innervate the OFC, a target that was densely innervated by KOR-lacking afferents. Lastly, KOR activation inhibited BLA inputs to non-fast spiking SOM neurons but not fast-spiking PV-positive interneurons, suggesting that KOR-lacking and KOR-expressing BLA inputs may differentially engage inhibitory mPFC microcircuits. Within mPFC circuits, multi-plexed encoding of information may in part be defined by projection patterns, molecular signatures, and/or localization of neurons across cortical layers, factors that are in large part inter-related (Otis et al., 2017; Ye et al., 2016; Condylis et al., 2022; Bugeon et al., 2022). Within the mPFC, Dyn and KOR-expressing neurons are found in different layers, raising the possibility that these different populations of neurons may differentially contribute to processing of limbic information and executive function. Heterogenous expression of KORs within cortico-cortical and amygdalo-cortical networks provides yet another mechanism by which the Dyn / KOR system may fine-tune the activity of specific mPFC neuronal ensembles to direct behavioral selection and executive control by the mPFC. Work from our laboratory demonstrates that mPFC Dyn cells are activated by threats and threat-predictive cues and release Dyn to change mPFC network states (Wang et al. 2022). Understanding whether Dyn and KOR-positive neurons and afferents pertain to specific ensembles of cells whose activity, connectivity, cell-type, and/or molecular identity is critical for valence processing, cognition, perception, and/or processing of general behavioral states is essential for understanding how the Dyn / KOR system shapes mPFC processing.

In addition to directly inhibiting excitatory drive arising from KOR-positive afferent inputs, Dyn acts via multiple mechanisms to suppress afferent-driven polysynaptic inhibition. Interestingly, KOR-mediated disinhibition occurs in a pathway-independent manner owing to actions on mPFC circuit elements downstream of extrinsic afferent inputs, including inhibition of excitatory KOR-expressing cells that recruit feedback inhibition and direct inhibition of GABA release from subsets of inhibitory interneurons that express KOR. Moreover, Dyn signaling decreases the probability that KOR-positive afferents will recruit SST-mediated feedforward mechanisms that would otherwise shunt incoming afferent inputs, such as those from the VH or the thalamus (Joffe et al., 2022; Tejeda and O’Donnell, 2014). KOR-mediated disinhibition of mPFC circuits is consistent with previous *in-vivo* microdialysis findings demonstrating that KOR activation inhibits the ability of a glutamate reuptake blocker to enhance extracellular GABA levels more potently than rises in extracellular glutamate (Tejeda *et al*., 2013). Dyn also inhibits sIPSCs onto principal neurons in the insular cortex (Pina et al., 2020), suggesting that the Dyn / KOR system may disinhibit other cortical circuits beyond the mPFC. In humans, KOR activation with synthetic KOR agonists or Salvinorin A drives potent psychotomimetic effects. Since thought and neurodevelopmental disorders associated with psychosis (e.g. schizophrenia) have been hypothesized to be impacted by deficits in PFC inhibitory circuits, this raises the intriguing possibility that the psychotomimetic effects of KOR agonists may be driven by cortical disinhibition.

Dyn acting via KORs may not just inhibit or disinhibit mPFC circuits, but rather may operate to shift which specific afferent inputs control mPFC circuits. KOR-mediated inhibition of BLA inputs is not overcome by high frequency activity (Fig. 8; Tejeda *et al*., 2015), suggesting that this system may not act as a high pass filter at these synapses, but rather turn down the gain of KOR-sensitive synapses. Similar to KORs, DA D1 receptor inhibition is sustained throughout on-going mPFC afferent activity (Burke et al., 2018), suggesting that sub-classes of GPCRs act to decrease the gain of mPFC networks. Via presynaptic KOR activation, Dyn may be decreasing the probability that glutamate released from KOR-expressing inputs will efficiently depolarize post-synaptic compartments and limit engagement of factors that promote synaptic plasticity and/or dendritic-somatic coupling. However, the disinhibitory effects of Dyn/KOR signaling enhances the ability of KOR-negative afferent inputs, such as the VH, that may be active at the time of Dyn release to control mPFC neuronal ensembles. Further, Dyn / KOR signaling may bias sub-networks of inhibitory neurons. Feedforward inhibition carried by SOM neurons is impacted by the Dyn / KOR system at two levels; 1) via presynaptic KORs that inhibit excitatory inputs onto SOM neurons and 2) direct inhibition of GABA release from KOR-positive SOM interneurons onto principal neurons. In contrast, PV-mediated feedforward inhibition is only subject to direct inhibition of GABA release from subsets of KOR-positive PV interneurons, since KOR activation does not modify excitatory inputs onto PV neurons. Differential modulation of sub-networks of mPFC interneurons has direct implications in cortical processing as PV- and SOM mediated feedforward inhibition of principal neurons diminishes and is enhanced with sustained afferent input activity, respectively (McGarry and Carter, 2016). Since SOM neurons innervate dendritic fields of principal neurons and exert increasing control of pyramidal neuron dendrites with on-going afferent input activity, Dyn-mediated disengagement of SST interneurons may amplify the spatiotemporal window by which excitatory inputs in dendrites may summate and trigger events critical for synaptic plasticity and excitation-spiking coupling (Palmer et al., 2014; Larkum et al., 2022; Lovett-Barron et al., 2012). Lastly, KORs are also expressed on presynaptic DA terminals in the mPFC where they directly inhibit DA release (Tejeda *et al*., 2013). DA produces complex effects of mPFC circuit function via interactions with various DA receptors that are in part segregated between cells in a layer specific manner (Anastasiades and Carter, 2021; Burke *et al*., 2018; Arnsten et al., 2015; Tseng and O’Donnell, 2004). Thus, in conjunction with regulation of excitation and inhibition balance, Dyn / KOR regulation of DA may further limit engagement of ensembles of mPFC neurons whose activity or synaptic integration is gated by DA receptor activation. Thus, Dyn release during motivationally-charged behaviors may influence mPFC processing by KORs expressed selectively within mPFC circuits and afferent inputs.

Dysregulated PFC Dyn / KOR expression and/or binding is observed with low social status, with childhood trauma, chronic exposure to alcohol, opioids, and/or life-long cannabis exposure, and has been implicated in mediating cognitive deficits during alcohol and opioid withdrawal (Wei et al., 2022; Abraham et al., 2021; Bazov et al., 2013; Bazov et al., 2018; Peckys and Hurd, 2001; Lutz et al., 2018). Furthermore, KOR agonists produce psychotomimetic effects in humans and animal models. These studies converge on the hypothesis that the aforementioned risk factors that dysregulate the PFC Dyn / KOR system may contribute to the development and/or maintenance of perceptual distortions, hallucinations, and/or disordered cognitive processing associated with a plethora of psychiatric disorders. Further, dysfunction in excitation/inhibition balance in the mPFC is implicated in these disorders, raising the possibility that dysregulation of the mPFC Dyn / KOR system and it’s control over excitation / inhibition balance may contribute to mPFC circuit deficits in psychiatric disorders. Future research aimed at identifying how Dyn / KOR signaling within specific mPFC circuit elements shapes mPFC processing of motivation, cognition and perception is warranted to address this knowledge gap and to elucidate novel therapeutic targets to ameliorate PFC deficits in psychiatric disorders.

## Figure Legends

**Fig. S1.**
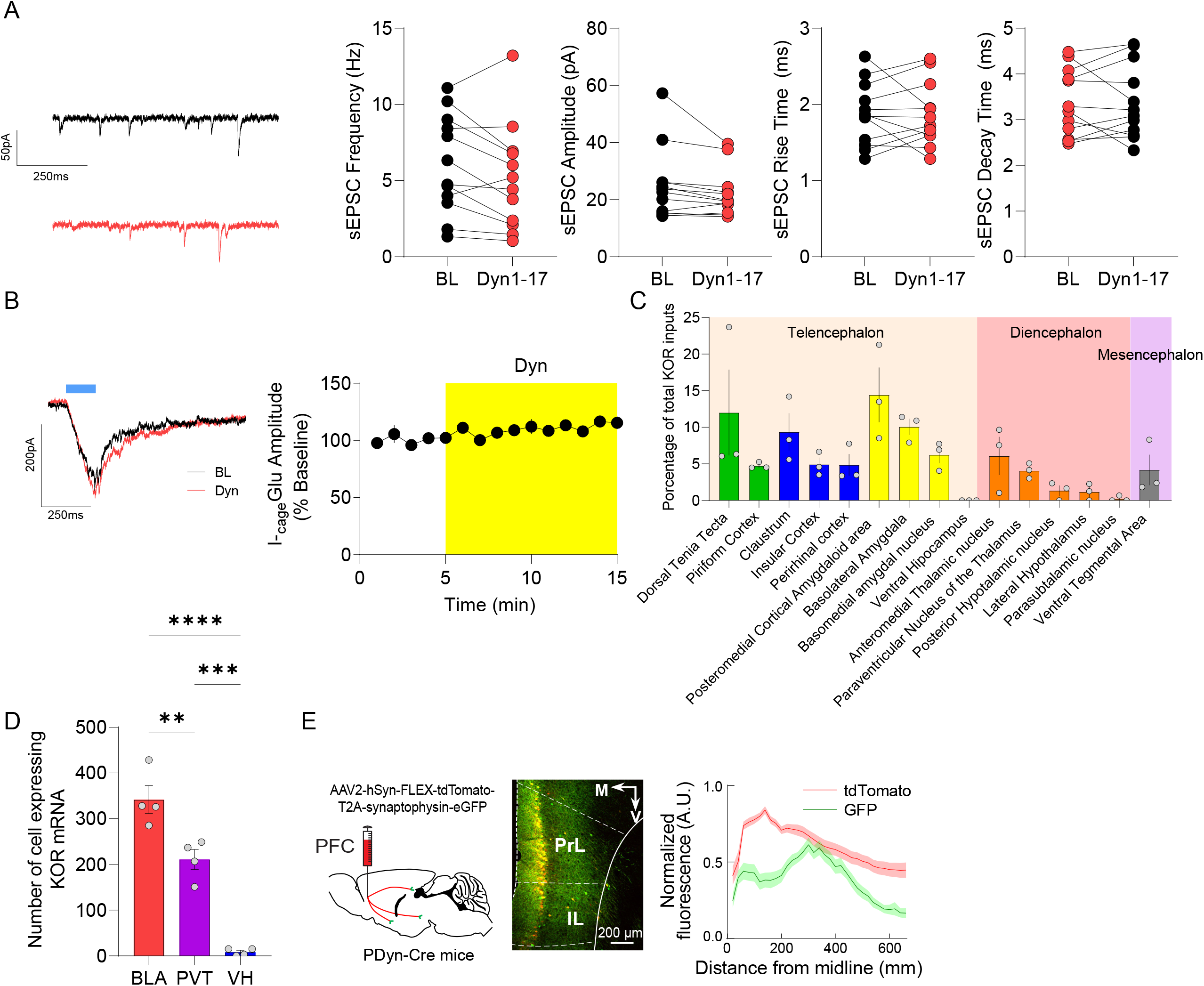
A. Dyn did not modulate sEPSC frequency or amplitude in mPFC cells, baseline (black), or Dyn (red). sEPSC frequency, amplitude, rise time, and decay time at baseline and during Dyn application (n = 12 cells, frequency, paired t-test, t_(11)_ = 1.97, *p* = 0.0735; amplitude, t_(11)_ = 1.98, *p* = 0.0729; rise time, t_(11)_ = 0.04, *p* = 0.9635; decay time, t_(11)_ = 0.008, *p* = 0.9936). B. Representative EPSCs recorded in mPFC pyramidal neurons evoked by MNI-Glutamate uncaging during baseline (black) and after Dyn (red). Dyn did not have an effect on the amplitude (expressed as a percentage of baseline) of the EPSC evoked by NMI-Glutamate in the mPFC (n = 6 cells). C. Percentage of KOR-expressing cells that project to PFC (n = 3 mice, ANOVA, F_(14, 30)_ = 3.83, *p* = 0.001). D. Number of cells expressing KOR mRNA in the PVT, BLA, and VH (one-way ANOVA, F_(2, 9)_ = 59.52, *p* < 0.0001, **p = 0.0055, ***p = 0.0003, ****p < 0.0001). E. Schematic and representative image of AAV-hSyn-FLEX-TdTomato-T2A-Synapsin-eGFP expression in the mPFC of PDyn-Cre mice. Normalized distribution of PFC Dyn cell arborization (tdTomato) and putative PFC Dyn synapses (eGFP lacking tdTomato) across layers of the PFC (n = 5 mice).

**Fig. S2.**
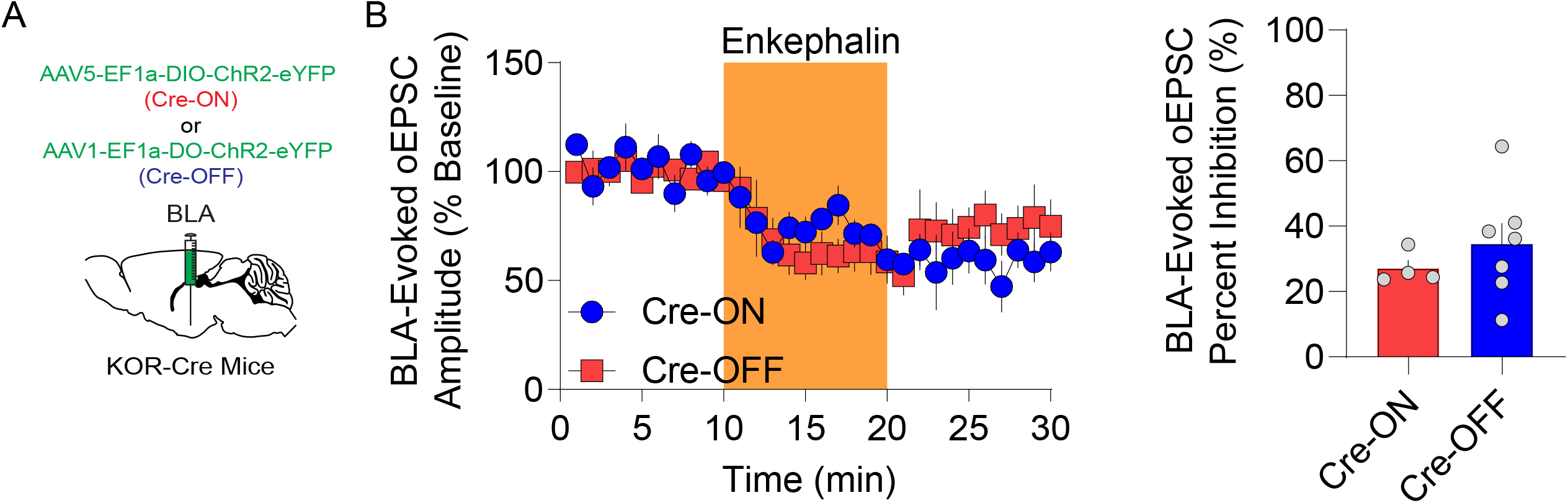
A. KOR-Cre mice were injected bilaterally with AAV5-EF1α-DIO-ChR2-eYFP (Cre-ON) or AAV1-EF1α-FLOX-ChR2-eYFP (Cre-OFF) into the BLA. B. Time course of enkephalin (1 µM) inhibition of oEPSC amplitudes in mPFC pyramidal neurons in animals expressing in KOR-containing (Cre-ON) (n = 4 cells) or KOR-lacking (Cre-OFF) (n = 7 cells) ChR2-eYFP in the BLA (Time course Two-way ANOVA; Time x virus interaction; F_(24, 168)_ = 0.90, *p* = 0.6058). Percentage of oEPSC inhibition by enkephalin application between KOR-containing (Cre-ON) and KOR-lacking (Cre-OFF) ChR2-eYFP in the BLA (t_(9)_= 0.85, p = 0.4139).

**Fig S3.**
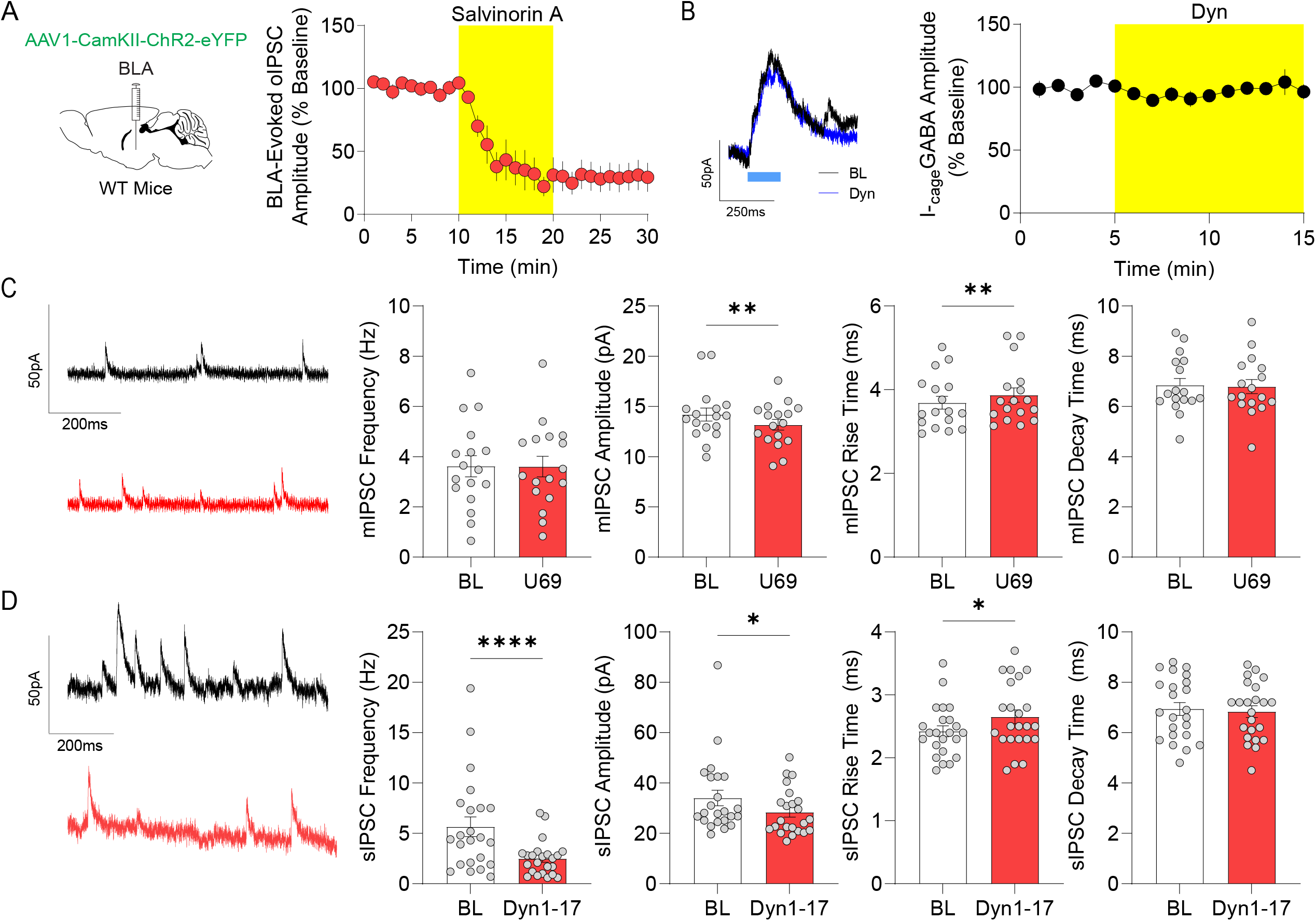
A. WT mice were injected bilaterally with AAV1-CaMKII-ChR2-eYFP in BLA of WT mice. Time course of the effect of KOR activation with Salvinorin A (1 µM) on the amplitude of oIPSCs in the mPFC of animals expressing ChR2-eYFP in the BLA (n = 6 cells). B. Representative oIPSCs evoked by Rubi-GABA uncaging in mPFC neurons. GABA uncaging-evoked IPSCs during baseline (black) and after Dyn1-17 (blue) are shown. Dyn did not have an effect on the amplitude of oIPSCs evoked by Cage-GABA in PFC (n = 6 cells). C. Representative mIPSC traces during baseline (black) or U69,593 application (red). Comparison of mIPSC frequency, amplitude, rise time, and decay time between baseline and U69,593 application (n = 17 cells, frequency, t_(16)_= 0.08, p = 0.9360; amplitude, t_(16)_ = 3.22, *p* = 0.0053; rise time, t_(16)_ = 3.51, *p* = 0.0029; decay time, t_(16)_= 0.38, *p* = 0.7020). D. Representative sIPSC traces of baseline (black) or Dyn1-17 (red). Comparison of sIPSC frequency, amplitude, rise time, and decay time between baseline and Dyn (n = 23 cells, frequency, t_(22)_ = 4.92, *p* < 0.0001; amplitude, t_(22)_ = 2.72, *p* = 0.0123; rise time, t_(22)_ = 2.24, *p* = 0.0354; decay time, t_(22)_ = 0.74, *p* = 0.4659).

**SF4.**
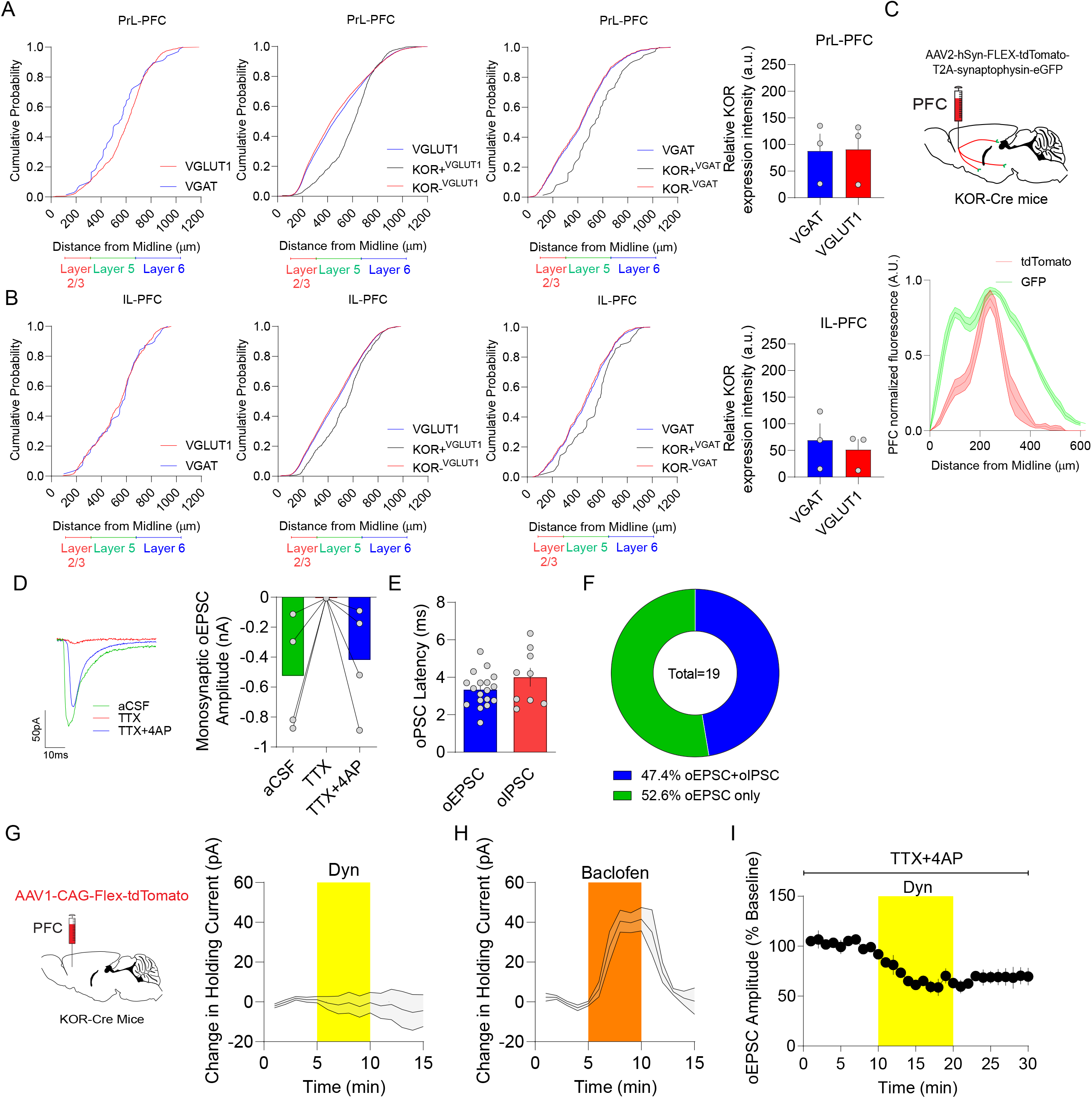
A. Cumulative probability of VGAT (blue) and VGLUT1 (red) mRNA across PrL-PFC layers. Cumulative probability of KOR mRNA expression with (black) and without (red) VGLUT1 across PrL-PFC layers. Cumulative probability of KOR mRNA expression with (black) and without (red) VGAT across PrL-PFC layers. Relative KOR mRNA expression in cells containing VGAT and VGLUT1 mRNA in PrL-PFC. Data are mean ± SEM (n = 3 mice, t_(4)_= 0.06, *p* = 0.9498). B. Cumulative probability of VGAT (blue) and VGLUT (red) mRNA across IL-PFC layers. Cumulative probability of KOR mRNA expression with (black) and without (red) VGLUT1 across IL-PFC layers. Cumulative probability of KOR mRNA expression with (black) and without (red) VGAT across IL-PFC layers. Relative KOR mRNA expression in cells containing VGAT and VGLUT1 mRNA in IL-PFC. Data are mean ± SEM (n = 3 mice, t_(4)_= 0.48, *p* = 0.6524). C. Schematic depicting AAV-hSyn-FLEX-TdTomato-T2A-Synapsin-eGFP expression in the mPFC of KOR-Cre mice. Normalized distribution of mPFC KOR cell arborization (tdTomato) and putative KOR synapses within the mPFC (eGFP signal lacking tdTomato) across mPFC layers (n = 4 mice; Two-way ANOVA; Fluorophore x distance interaction; F_(31, 486)_ = 7.43, *p* < 0.0001). D. Representative traces of oEPSC from KOR cells in the PFC that make excitatory monosynaptic connections. Mean monosynaptic oEPSC amplitude in the presence of aCSF, TTX and TTX+4AP. Data are mean ± SEM (n = 4 cells; t_(25)_=1.46, *p*=0.1563). E. Mean monosynaptic oEPSC and oIPSC latency consistent with direct monosynaptic excitatory and inhibitory connections. Data are mean ± SEM (n = 19 cells). F. Pie chart showing the proportion of evoked responses consisting of oEPSC+IPSC and oEPSC from KOR-positive neurons. G. KOR-Cre mice were injected bilaterally with AAV1-CAG-Flex-tdTomato into PFC. Time course of the effect of KOR activation with Dyn on change in holding current on mPFC KOR neurons (n = 10 cells). H. Time course of the effect of GABA-B activation with baclofen (1 µM) on change in holding current in mPFC principal neurons (n = 10 cells). I. The amplitude of evoked monosynaptic oEPSCs from KOR-positive neurons onto mPFC pyramidal neurons are inhibited by Dyn (n = 4 cells).

**SF5.**
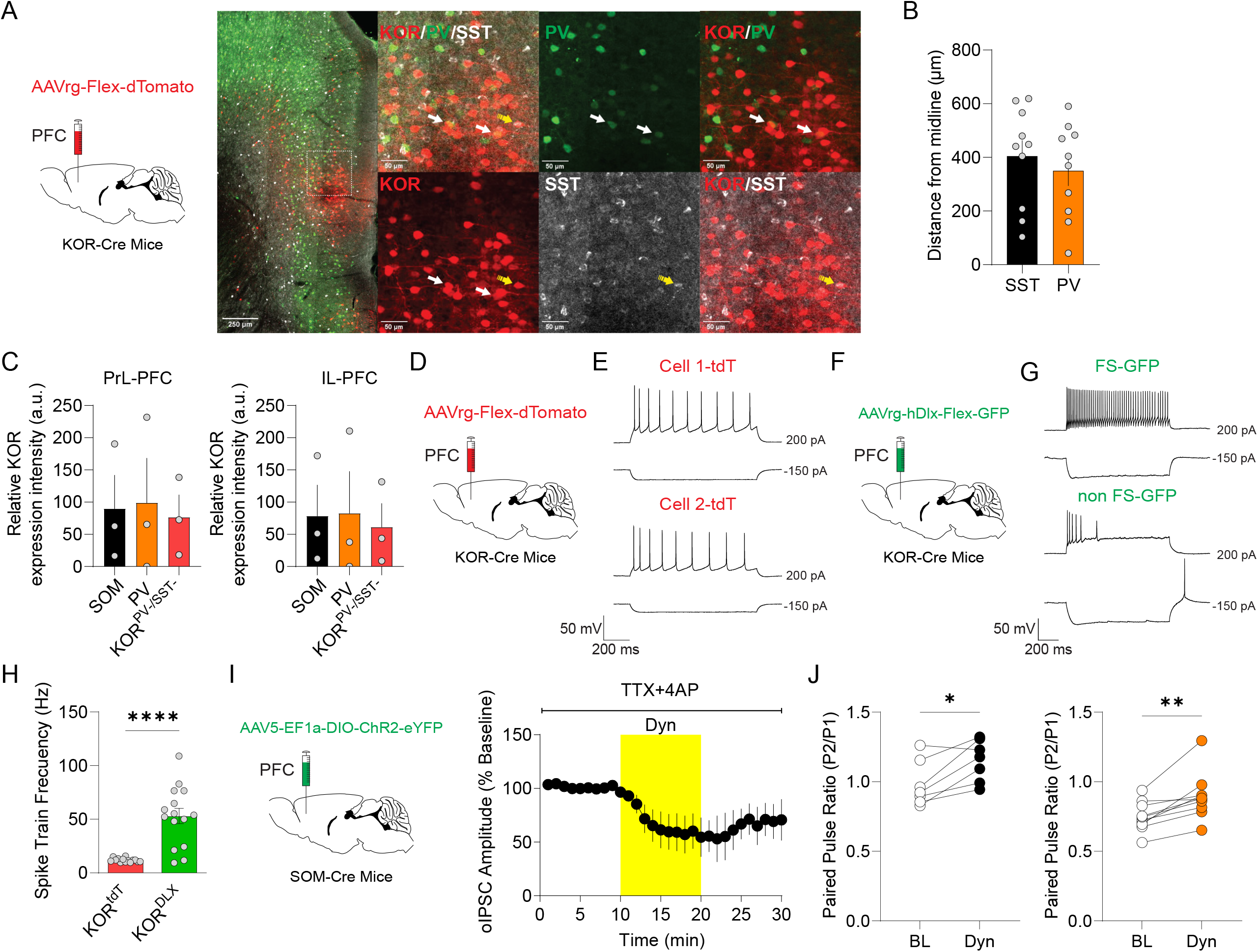
A. KOR-Cre mice were injected unilaterally with AAVrg-FLEX-tdTomato into the mPFC. Representative images show colocalization of tdTomato-labeled mPFC KOR-expressing cells with PV-(green) and SOM-immuroreacitivity in a sparse population of tdTomato-positive cells (white; n = 3 mice). B. Relative distance from midline of KOR-positive tdTomato cells colocalized with PV or SOM-immunoreactivity, which is similar to the distribution KOR mRNA-positive PV and SOM-mRNA positive cells as assessed by *in-situ* hybridization in WT mice (t_(18)_= 0.67, *p* = 0.5104). C. Relative KOR mRNA expression in cells containing or lacking SOM and PV mRNA in PrL- and IL-mPFC (n = 3 mice, PrL-mPFC ANOVA, F_(2, 6)_ = 0.044, *p* = 0.9565; IL-mPFC ANOVA, F_(2, 6)_ = 0.049, *p* = 0.9524). D. Schematic showing bilateral injection of AAV1-CAG-Flex-tdTomato into the mPFC of KOR-Cre mice to label all KOR-positive neurons. E. Representative traces of intrinsic excitability at different current steps of two tdTomato positive cells, consistent with regular firing pyramidal neurons. F. KOR-Cre mice were injected bilaterally with AAVrg-hDlx-Flex-GFP into the mPFC to label KOR-positive inhibitory interneurons. G. Representative traces at different current steps showing intrinsic properties consistent with a fast-spiking interneuron (top) and a non-fast spiking, non-regular spiking interneuron (bottom). H. Comparison of spike train frequency between KOR-excitatory and inhibitory cells (KOR-tdT n = 14 cells; KOR-Dlx n = 19 cells, t_(27)_= 5.52, p < 0.0001). I. Schematic depicting SOM-Cre mice injected bilaterally with AAV5-EF1α-DIO-ChR2-eYFP into the mPFC. Time course of monosynaptic oIPSCs in the presence of TTX/4-AP demonstrating the inhibitory effect Dyn (1 µM) application from animals expressing ChR2-eYFP in SOM-positive mPFC neurons (n = 4 cells). J. Comparison of the effect of KOR activation by Dyn on the paired-pulse ratio of oIPSC evoked from SOM-positive (black, t_(6)_= 3.46, *p* = 0.0134) and PV-positive (orange, t_(9)_= 4.37, *p* = 0.0018) neurons in the mPFC.

**SF7.**
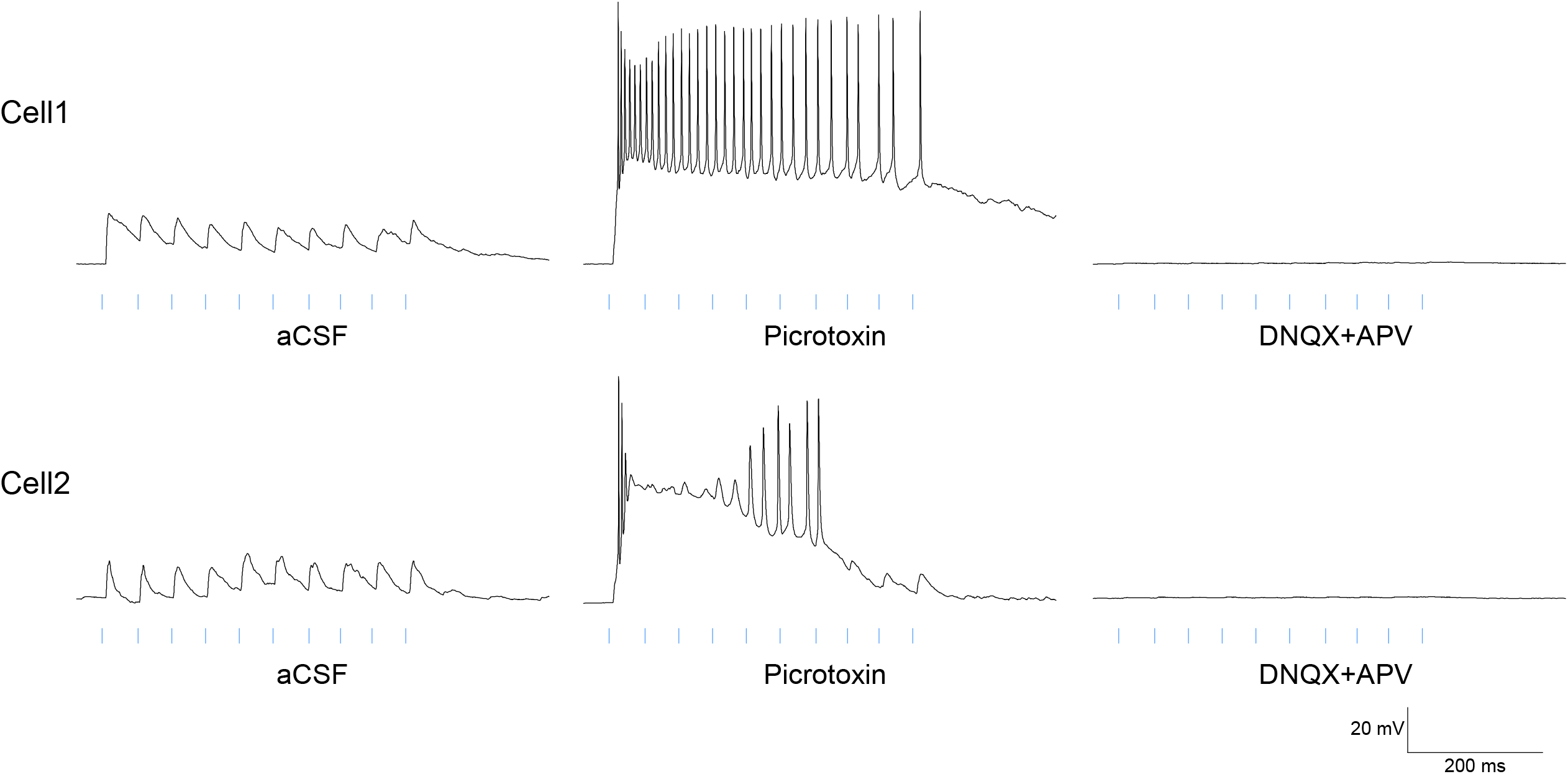
Representative traces of two neurons demonstrating that synaptically driven action potential firing by train stimulation of VH inputs (10 pulses at 20 Hz) is under the control of GABAergic inhibition. Evoked spiking during baseline, in the presence of picrotoxin, and after DNQX+APV application. Blue lines indicate timing of LED pulses

## Methods

### Animals

Adult (> postnatal day 60) male and female C57/Bl6J WT, prodynorphin-Cre (PDyn-iCre), somatostatin-Cre (SOM-Cre), parvalbumin-Cre (PV-Cre), kappa opioid receptor-Cre (KOR-Cre), and kappa opioid receptor-loxP (KOR-loxP) mice were used. Mice were housed in humidity and temperature-controlled vivariums using a reverse light cycle with lights off at 7:00 hours and lights on at 19:00 hours. Animals had ad-libitum access to standard laboratory chow and water. Mice were obtained from Jackson Laboratories. All procedures were approved by the National Institute of Mental Health Animal Care and Use Committee.

### Viral Injections

Mice were anesthetized by ketamine/xylazine cocktail and head-fixed in a stereotaxic frame. For viral injections experiments, subjects were injected with 300-500 nL of virus bilaterally or unilaterally depending on the experiment. The coordinates utilized for the medial prefrontal cortex (mPFC, from bregma: A/P +1.70, M/L +0.3, D/V 2.40 mm), ventral hippocampus (VH, from bregma: A/P 3.27, M/L 3.15, D/V 4.95), paraventricular thalamus (PVT, from bregma: A/P +0.06, M/L -1.58, D/V 3.2 with a 6° degree angle) and basolateral amygdala (BLA, from bregma: A/P -1.65, M/L 3.25, D/V 5.00). Experiments were then performed approximately 4-8 weeks after injection (1 week for retrobeads experiments) to allow sufficient time for expression and trafficking. Acute slices were prepared from most subjects, except for RNAscope or immunohistochemistry experiments where brains were freshly extracted or subjects were perfused, respectively.

### Immunohistochemistry

Mice were transcardially perfused with phosphate-buffered saline (PBS) followed by 4% paraformaldehyde. Brains were post-fixed in 4% paraformaldehyde for 4 hr and subsequently transferred to PBS. Brain slices (50 µm) containing the mPFC were obtained using a vibratome (Leica). Slices were washed 3 times in PBS, blocked in PBS containing 0.2% Triton-X and 4% bovine serum albumin for 2 hrs. Slices were then incubated with rabbit anti-SOM (1:200) and anti-PV (1:1000) primary antibodies in a blocking solution overnight. The following day sections were washed for 5 min in PBS 3 times. Brain slices were then incubated with donkey anti-mouse (488 nm; 1:500 dilution) and donkey anti-rat secondary (647 nm; 1:200 dilution) for 2 hrs. Slices were then washed for 5 min, 4 times, and mounted. Confocal images were acquired using a confocal microscope (Zeiss 780 LSM). For quantification of synaptophysin and BLA inputs to the mPFC, the ROI image was transformed into an image text file to quantify fluorescence in 20 µm bins, and data were normalized to the maximum value of fluorescence.

### Retrobead retrograde tracing

C57/Bl6J WT mice were anesthetized and secured on a stereotaxic frame, as described above. Green XI retrobeads (Lumafluor) were unilaterally injected into the mPFC. Retrobeads (150-300 nl) were injected into the mPFC (AP: +1.7 mm; ML: ±0.3 mm; DV: -2.4 mm) and the injector was left in place for 8 min after injection. Brains were removed for RNAscope in situ hybridization 6-7 days after surgery.

### RNAscope in situ hybridization (ISH)

Brains were rapidly dissected, and flash frozen with isopentane cooled with dry ice for 20 sec and stored at -80°C until sectioning for ISH. Brain slices (16 µm) containing the mPFC, PVT, BLA, and VH were obtained utilizing a Leica CM 3050S cryostat at -20°C and were mounted directly onto microscope slides cleaned with RNAzap, to prevent mRNA degradation. Slides containing ISH sections were stored at -80°C until ISH processing. RNAscope ISH was conducted according to the Advanced Cell Diagnostics user manual and as previously reported (Tejeda et al., 2013, 2017). Briefly, slides were fixed in 10% neutral buffered formalin for 20 min at 4°C. Slides were subsequently washed twice for 1 min with PBS, before dehydration with 50% ethanol (5 min), 70% ethanol (5 min), and 100% ethanol (5 min). Slides were incubated in 100% ethanol at -20°C. The following day, slides were dried at room temperature (RT) for 10 min. A hydrophobic barrier was drawn around the sections using a hydrophobic pen and allowed to dry for 10-15 min at RT. Sections were then incubated with Protease Pretreat-4 solution for 20 min at RT. Slides were washed with ddH2O (2 × 1 min), before being incubated with the appropriate probes for 2 hr at 40°C in the HybEZ oven (Advanced Cell Diagnostics). Probes used were purchased from Advanced Cell Diagnostics and are as follows: Mm-Oprk1-C1 (nucleotide target region 256-1457; Accession number NM_001204371.1), Mm-Pvalb-C2 (nucleotide target region 2-885; Accession number NM_013645.3), Mm-Sst-C3 (nucleotide target region 18-407; Accession number NM_009215.1), Mm-Slc17a7-C2 (nucleotide target region 621-1021; Accession number NM_182993.2), Mm-Slc32a1-C3 (nucleotide target region 894-2037; Accession number NM_009508.2), Mm-Pdyn-C2 (nucleotide target region 33-700; Accession number NM_0188863.3). Slides were washed in wash buffer twice for 2 min, prior to being incubated with Amplification 1 buffer at 40°C in the HybEZ oven for 30 min. Slides were subsequently washed in wash buffer twice for 2 min, then incubated with Amplification 2 buffer at 40°C in the HybEZ oven for 15 min. Slides were washed in wash buffer twice for 2 min, prior to being incubated with Amplification 3 buffer at 40°C in the HybEZ oven for 30 min. Slides were subsequently washed in wash buffer twice for 2 min and incubated with Amplification 4-Alt A buffer at 40°C in the HybEZ oven for 15 min. Slides were washed in wash buffer twice for 2 min. DAPI solution was applied to sections at RT for 20 sec. Finally, slides were coverslipped and stored at 4°C until imaging on a confocal microscope (Olympus). A custom pipeline developed in Cell Profiler was used for RNAscope ISH analysis (Erben and Buonanno, 2019).

### Acute brain slice preparation

Mice were anesthetized with euthanasia (NIH Veterinarian Services) and subsequently decapitated. Brains were removed rapidly and placed in ice-cold NMDG-based cutting solution containing (in mM): 92 NMDG, 20 HEPES, 25 glucose, 30 NaHCO_3_, 2.5 KCl, 1.2 NaPO_4_ saturated, 10 Mg-sulfate, and 0.5 CaCl_2_ with 95% O_2_/5% CO_2_ with an osmolarity of 303-306 mOsm (Wescorp). The brain was rapidly blocked, dried on filter paper, and glued to a platform containing ice-cold NMDG-based cutting solution in a chamber placed within a Leica VT1200 Vibratome. Coronal slices (250 µm thick) containing the mPFC were cut at a speed of 0.07 mm/s. Following slicing, sections were incubated in a chamber containing NMDG-based cutting solution for 5-10 min at 34°C. Slices were subsequently transferred to a chamber filled with a modified holding aCSF saturated with 95% O_2_/5% CO_2_ containing (in mM): 92 NaCl, 20 HEPES, 25 glucose, 30 NaHCO_3_, 2.5 KCl, 1.2 NaPO_4_, 1 mM Mg-sulfate and 2 mM CaCl_2_ (303-306 mOsm) at room temperature for at least 1 hr. Slices remained in this solution until being transferred to the recording chamber.

### Ex-vivo whole cell electrophysiology

Whole-cell patch-clamp electrophysiology studies were performed as previously described ((Tejeda et al., 2017); (Pignatelli et al., 2021)). Cells were visualized using IR-DIC optics on an inverted Olympus BX5iWI microscope. For recordings, the recording chamber was perfused with a pump (World Precision Instruments) at a flow rate of 1.5-2.0 ml per minute with aCSF containing (in mM): 126 NaCl, 2.5 KCl, 1.4 NaH_2_PO_4_, 1.2 MgCl_2_, 2.4 CaCl_2_, 25 NaHCO_3_, and 11 glucose (303-305 mOsm). For biophysically isolated oEPSCa and oIPSCs cells were held at - 55mV and +10mV respectively. Monosynaptic oEPSCs and oIPSCs evoked by KOR^+^ cell stimulation were isolated with TTX (1 µM) and 4-AP (50 µM). For whole-cell recordings of intrinsic excitability, we utilized glass microelectrodes (3-5 MΩ) containing (in mM): 135 K-gluconate, 10 HEPES, 4 KCl, 4 Mg-ATP, and 0.3 Na-GTP. For oEPSCs and oIPSCs, excitation/inhibition balance, Rubi-GABA evoke IPSCs, NMI-glutamate evoked EPSC and NMI-glutamate evoked IPSCs at -55mV and +10 mV we utilized glass microelectrodes (3-5 MΩ) containing (in mM): 117 cesium methanesulfonate, 20 HEPES, 0.4 EGTA, 2.8 NaCl, 5 TEA-Cl, 4 Mg-ATP, 0.4 Na-GTP (280-285 mOsm). ChR2-negative cells were identified by the lack of ChR2 currents evoked by blue light stimulation. ChR2 currents were characterized by sustained, steady-state currents in response to 100 ms blue light stimulation with an onset at the start of the laser pulse. For paired pulse ratio quantification, light evoked currents were recorded in response to two light pulses with a 50 ms interstimulus interval. To determine the effects of KOR activation, a ten-minute baseline was collected, and Dyn A 1-17or Salvinorin A was subsequently bath applied for 10 min and after the drug was washed for ten minutes. The last five minutes of baseline and the last five minutes of the drug were used for quantification. To determine the relative engagement of polysynaptic and monosynaptic excitation and inhibition by mPFC KOR-positive neurons, recordings were made in aCSF or isolating monosynaptic current with TTX and 4AP, respectively. oEPSCs (0.1 Hz) were recorded at -55 mV until the response was stable. Voltage was then gradually ramped up to +10 mV to biophysically isolate IPSCs. Neurons were voltage-clamped utilizing a Multiclamp 700B amplifier (Molecular Devices). Data were filtered at 2 kHz and digitized at 20 kHz using a 1440A Digidata Digitizer (Molecular Devices). Series-resistance (<20 MΩ) was monitored using a -5 mV voltage step. Cells with >20% change in series resistance were discarded from further analysis. For intrinsic excitability, after membrane rupture in voltage clamp, cells were switched to the current clamp configuration without holding current injection. Miniature EPSCs (mEPSCs} were recorded in the presence of 1 µM TTX and picrotoxin (100 µM). Miniature IPSCs (mIPSCs) were collected in the presence of TTX (1 µM), DNQX (10 µM), and D-AP5 (50 µM). For sEPSCs, sIPSCs, mEPSCs and mIPSCs a ten-minute baseline was collected and U69,593 or Dyn A 1-17 was subsequently bath applied. The last five minutes of baseline and the last five minutes of drug were used for quantification and were counted manually utilizing Minianalysis software (Synaptosoft). Change in holding current by Dyn1-17 was evaluated at -60 mV, a positive control was used Baclofen 10µM. MNI-glutamate was bath applied at 50 µM in 10 ml of ACSF. Experiments at -55 mV and +10 mV were performed to isolate uEPSC and uIPSC, respectively. DNQX+AP5 was added to demonstrate that glutamate uncaging-evoked IPSCs were driven by glutamate receptor activation. Glutamate uncaging was achieved using a 150 ms pulse of n before the drug trials. Intrinsic excitability was assessed by applying hyperpolarizing and depolarizing current steps (25 pA steps; 1-sec duration) and measuring the change in voltage and action potential firing. For experiments determining the effects of Dyn A 1-17 on synaptically-driven mPFC neuron spiking, whole-cell recordings were performed using a potassium gluconate-based internal solution. Optogenetic synaptically-driven spiking was evoked by a 10 pulse 20 Hz train stimulation delivered every 20 sec. In some experiments, stimulation intensity was adjusted so that cells exhibited spikes evoked by optogenetic train stimulation or failed to evoke spikes in the case of subthreshold experiments. A five-minute baseline was collected, and Dyn A 1-17 was subsequently bath applied. The last two minutes of baseline and the last two minutes of drug were used for quantification.

### Two-Photon Imaging

Imaging was performed by using an upright FVMPE-RS multiphoton microscope (Olympus) equipped with an InSight DS Dual-OL fs-laser system (Spectra-Physics) and equipped with 40X, 0.8 NA water-immersion objective (Olympus). For 2PLSM, 940 nm light was used to excite the genetically-encoded calcium sensor jGCaMP7f. Reference frame scans were taken between each acquisition to correct for small spatial drift of the preparation over time. Optogenetic synaptically-driven spiking was evoked by a 10 pulse 20 Hz train stimulation delivered every 30 sec by a 635 nm laser directed at the slice containing the ROI being imaged with the microscope. The stimulation intensity used was approximately 60 mW. To measure Ca^2+^ signals, green fluorescence was collected during 15 Hz full field scan using a 512×512 resonant scanner. Ca^2+^ signals were quantified as changes in fluorescence relative to the baseline time. Images were analyzed off-line with ImageJ (NIH). Data are presented as the area under the curve in fluorescence with respect to the baseline period.

### Drugs

Drugs were dissolved in aCSF or water. Drugs were purchased from Sigma Aldrich, Tocris, or generously provided by the NIDA Drug Supply Program.

### Statistics

Statistics were computed using GraphPad Prism. Detailed statistics can be found in supplemental Table 1.

## Acknowledgements

This work was supported by the NIMH Intramural Research Program (ZIA MH002970-04), a NARSAD Young Investigator Award from the Brain and Behavior Research Foundation (HAT), and a NIH Center for Compulsive Behaviors Fellowship (HEY). The authors would like to thank Drs. Mario Penzo, Eastman Lewis, Andres Buonanno, Marco Pignatelli, and members of the Tejeda laboratory for their comments on the study and manuscript. The authors would like to thank Drs. Yogita Chudasama and Claire Le Pichon for generously sharing their cryostats for sample preparation. The authors would like to thank Drs. Ted Usdin and Sarah Williams of the NIMH Systems Neuroscience Imaging Resource.

